# Development and validation of an expanded antibody toolset that captures alpha-synuclein pathological diversity in Lewy body diseases

**DOI:** 10.1101/2022.05.26.493598

**Authors:** Melek Firat Altay, Senthil T. Kumar, Johannes Burtscher, Somanath Jagannath, Catherine Strand, Yasuo Miki, Laura Parkkinen, Janice L. Holton, Hilal A. Lashuel

## Abstract

The abnormal aggregation and accumulation of alpha-synuclein (aSyn) in the brain is a defining hallmark of synucleinopathies. Different aSyn conformations and post-translationally modified forms accumulate in pathological inclusions and vary in abundance across synucleinopathies. Relying on antibodies that have not been assessed for their ability to capture the diversity of aSyn species may not provide an accurate estimation of aSyn pathology in human brains or disease models. To address this challenge, we developed and characterised an expanded antibody panel that targets different sequences and post-translational modifications along the length of aSyn and recognises all three aSyn conformations (monomeric, oligomeric, fibrillar). Next, we profiled aSyn pathology across sporadic and familial Lewy body diseases (LBDs) and reveal heterogeneously modified aSyn pathologies rich in Serine 129 phosphorylation but also in Tyrosine 39 nitration and N- and C-terminal tyrosine phosphorylations, scattered to neurons and glia. We also show that aSyn may become hyperphosphorylated during the aggregation and inclusion maturation processes in neuronal and animal models of aSyn aggregation and spreading. The antibody validation pipeline we describe here paves the way for more systematic investigations of aSyn pathological diversity in the human brain and peripheral tissues, and in cellular and animal models of synucleinopathies.

## INTRODUCTION

Synucleinopathies are a group of neurodegenerative diseases that include Parkinson’s disease (PD), dementia with Lewy bodies (DLB) and multiple system atrophy (MSA) (Baba et al., 1998; Spillantini et al., 1998a, 1998b). Mounting evidence points to the aggregation and accumulation of the pre-synaptic protein alpha-synuclein (aSyn) as critical processes in the pathophysiology of synucleinopathies: 1) Fibrillar and aggregated forms of aSyn are enriched in the pathological inclusions of PD, DLB and MSA, including Lewy bodies (LBs), Lewy neurites (LNs), glial cytoplasmic inclusions (GCIs) and neuronal cytoplasmic inclusions (NCIs) (Baba et al., 1998; Spillantini et al., 1998b, 1998a); 2) mutations and duplications of the aSyn-encoding gene *SNCA* are shown to cause familial forms of PD (Appel-Cresswell et al., 2013; Chartier-Harlin et al., 2004; Kapasi et al., 2020; Kiely et al., 2013; Krueger et al., 1998; Lesage et al., 2013; Pasanen et al., 2014; Polymeropoulos et al., 1997, 1996; Proukakis et al., 2013; Singleton et al., 2003; Spira et al., 2001; Zarranz et al., 2004); and 3) inoculation of recombinant aSyn fibrils or PD and MSA brain-derived aSyn aggregates induces LB-like inclusion formation in cellular and animal models (Kumar et al., 2021; Luk et al., 2012, 2009; Mahul-Mellier et al., 2020, 2018; Tarutani et al., 2016; Volpicelli-Daley et al., 2014) and the spreading of LB-like intracellular pathology in brain regions and along the gut-brain axis (Arotcarena et al., 2020; Dehay and Bezard, 2019; Recasens et al., 2014; Rey et al., 2016, 2013).

Diverse forms of aSyn accumulate in brain inclusions associated with synucleinopathies (Anderson et al., 2006; Schweighauser et al., 2020; Shahmoradian et al., 2019), exhibiting variations in morphology (Kuusisto et al., 2003), biochemical composition (Leverenz et al., 2007; Wakabayashi et al., 2013; Xia et al., 2008), structure (Shahmoradian et al., 2019; Strohaeker et al., 2019) and distribution (Alafuzoff et al., 2009). This heterogeneity is influenced by factors such as the type of synucleinopathy, cell type, brain region, and patient clinical history (Figure 1A). Cryogenic electron microscopy (cryo-EM) studies have shown that aSyn can form fibrils with different morphologies and conformations *in vitro*, using full-length, truncated and modified recombinant and semi-synthetic aSyn monomers (Guerrero-Ferreira et al., 2019; Li et al., 2018) (Figure 1B). Similarly, fibrils isolated from PD, DLB and MSA brains also exhibit polymorphisms (Schweighauser et al., 2020; Yang et al., 2022).

**Figure 1:**
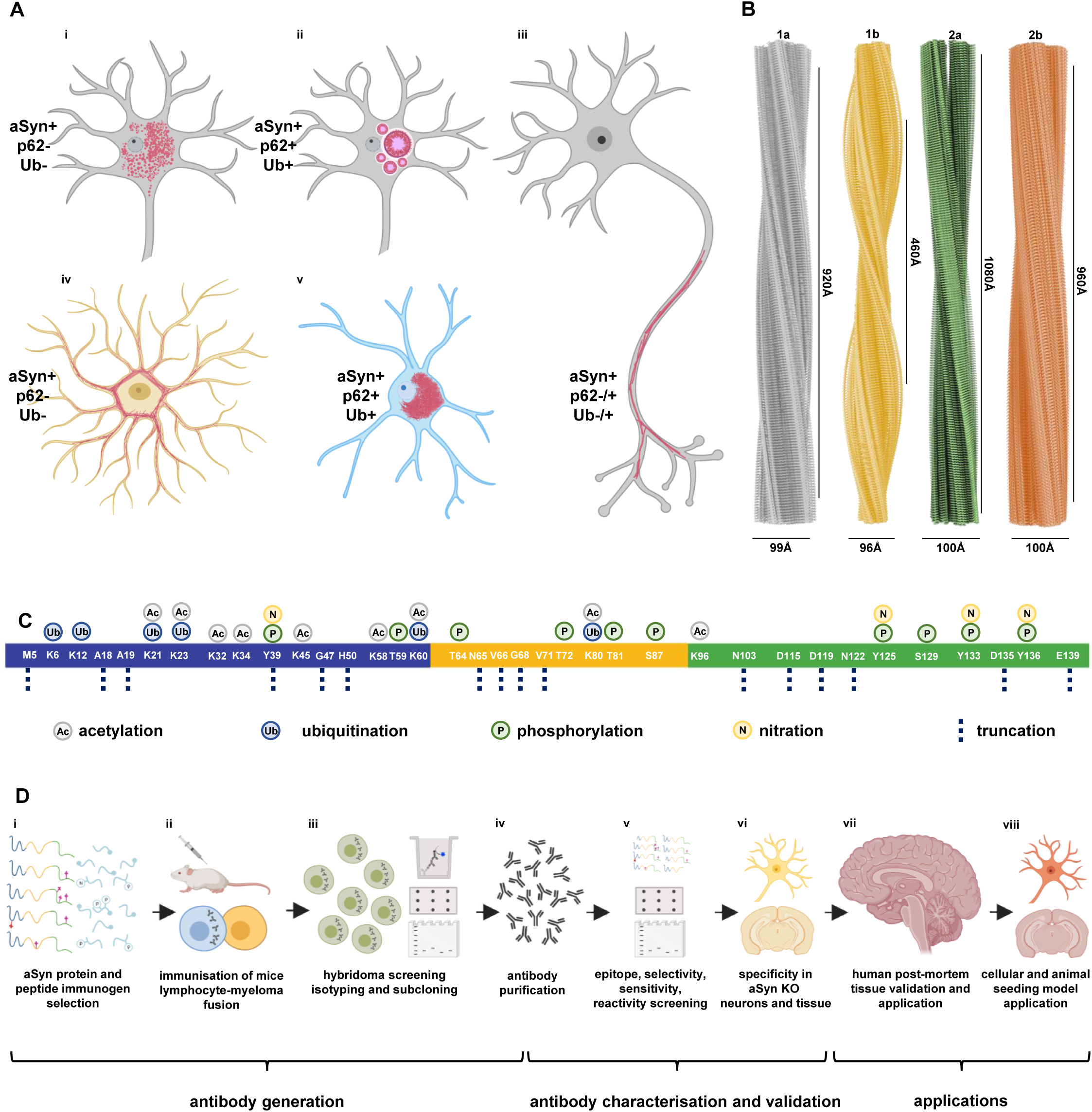
(A) Diversity of aSyn pathology in synucleinopathies with **(Ai)** granular/ punctate cytoplasmic inclusions in the neurons; **(Aii)** classical LBs in the neuronal soma; **(Aiii)** LNs in the neuronal processes; **(Aiv)** astrocytic aSyn accumulations; **(Av)** oligodendroglial cytoplasmic inclusions. These pathological structures show differences in their positivity to aggregation markers, including ubiquitin (Ub) and p62. Schematic created with BioRender.com (agreement no: *QW23G6FJ76*). **(B)** Cryo-EM three-dimensional reconstructions of the recombinant full-length aSyn PFFs to show the polymorphism of aSyn fibrils generated *in vitro* (Guerrero-Ferreira et al., 2019; Li et al., 2018). Four distinct polymorphs were identified based on the protofilament fold and inter-protofilament interfaces: Polymorph 1a ‘rod’ (PDB-6CU7, EMD-7618); polymorph 1b ‘twister’ (PDB-6CU8, EMD-7619); polymorph 2a (PDB-6SSX, EMD-10307); and polymorph 2b (PDB-6SST, EMD-10305). **(C)** aSyn PTMs identified in synucleinopathy brain tissues, which include acetylation, ubiquitination, phosphorylation, nitration and truncation across the whole sequence of the protein. **(D)** A schematic representation of the steps followed for the generation, characterisation, validation and application of aSyn antibodies. These involved (**Di**) antibody design via the selection of immunogens comprising of aSyn recombinant proteins and peptides; (**Dii**) immunisation of the mice followed by lymphocyte-myeloma fusion; (**Diii**) screening of the hybridomas via ELISA, DB and WB, isotyping and subcloning, and (**Div**) acquisition of purified antibodies. The antibodies were then (**Dv**) characterised using a library of aSyn recombinant proteins for their epitopes, conformational selectivity, sensitivity, specificity and reactivity via DB and WB. The antibody specificity was further validated on (**Dvi**) aSyn KO mouse primary hippocampal and cortical neurons, and in aSyn KO mouse tissue of amygdala. (**Dvii**) The antibodies were validated on human brain tissues of different LB disorders. (**Dviii**) The mouse aSyn-reactive antibodies were applied to neuronal seeding model and PFF-injected mouse brain tissues to profile the newly formed aggregates. Schematic created with BioRender.com (agreement no: *FN23G6E1SR*). aSyn = alpha-synuclein; DB = dot/slot blot; cryo-EM = cryogenic electron microscopy; ELISA = enzyme-linked immunoassay; KO = knockout; LB = Lewy body; LN = Lewy neurite; PFFs = pre-formed fibrils; PTM = post-translational modification; Ub = ubiquitin; WB = Western blot

The field of aSyn research has been largely relying on antibodies targeting different sequences, modifications or conformations of the protein to detect, quantify and monitor aSyn pathology in LBs, LNs, GCIs and NCIs (Covell et al., 2017; Dhillon et al., 2017; Duda et al., 2002; Fayyad et al., 2020; Giasson et al., 2000b; Henderson et al., 2020; Kovacs et al., 2012; Vaikath et al., 2015; Waxman et al., 2008). aSyn accumulating within the neuronal and glial inclusions is subjected to different types of post-translational modifications (PTMs) at multiple residues (Figure 1C), including phosphorylation, ubiquitination, nitration, acetylation and N- and C-terminal truncations. Of these modifications, aSyn phosphorylated at Serine 129 (aSyn pS129) has been described as the most common aSyn PTM in pathological aSyn aggregates, and aSyn pS129 levels have been reported to increase by multiple folds in PD, DLB and MSA brains (Anderson et al., 2006; Fujiwara et al., 2002). These observations have led to the emergence of aSyn pS129 as a key marker of aSyn pathology. Several antibodies against aSyn pS129 have been developed to assess aSyn pathology formation and spread in the central nervous system and peripheral tissues. However, the comprehensive coverage of aSyn pathology by these antibodies has not been thoroughly investigated.

Immunohistochemical and mass spectrometry studies on the distribution of aSyn species in LBs and GCIs revealed the presence of aSyn proteoforms cleaved at different residues within the C-terminal region of the protein (e.g. 1-115, 1-119, 1-122, 1-133, 1-135) (Anderson et al., 2006; Baba et al., 1998; Campbell et al., 2001; Dickson et al., 1999; Gai et al., 1999; Kellie et al., 2015; Lewis et al., 2010; Moors et al., 2019; Ohrfelt et al., 2011; Tofaris et al., 2003; Tong et al., 2010), and of other aSyn C-terminal modifications, including phosphorylation and/or nitration at tyrosine residues 125 (Y125), 133 (Y133) and 136 (Y136) (Chen et al., 2009; John E. Duda et al., 2000; Giasson et al., 2000a; Mahul-Mellier et al., 2014) (Figure 1C). Interestingly, the most abundant C-terminally truncated species, i.e. aSyn 1-119 and 1-122 (Anderson et al., 2006; Bhattacharjee et al., 2019; Kellie et al., 2015; Killinger et al., 2018; Ohrfelt et al., 2011), lack the epitope for aSyn pS129 antibodies. Furthermore, the occurrence of several modifications in close proximity to S129 may interfere with the immunoreactivity of several aSyn C-terminal or pS129 antibodies (Lashuel et al., 2022; Mahul-Mellier et al., 2018).

A large part of the immunohistochemistry and biochemistry studies on human brain samples, including the PD staging work by Braak and colleagues (Braak et al., 2001, 2003, 2002), has used antibodies with epitopes against the C-terminal region of aSyn. These commonly used antibodies are less likely to capture truncated forms of aSyn, aggregated but not yet phosphorylated aSyn, or aggregated aSyn bearing multiple C-terminal modifications (Anderson et al., 2006; Chen et al., 2009; Giasson et al., 2000a; Kiely et al., 2013; Mahul-Mellier et al., 2014). For example, astrocytic aSyn accumulations, composed of aSyn species that lack approximately the first and last 30 amino acids of the protein, are not revealed by aSyn C-terminal antibodies (Altay et al., 2022). Furthermore, the cellular environment influences the kinetics of aSyn aggregation, fibril polymorphism and PTMs (Peng et al., 2018). It is likely that the aSyn aggregates formed in the different synucleinopathy types, regions or disease stages, possess distinct biochemical and structural signatures that expose amino acid sequences and antibody epitopes differentially. Relying on a single antibody may underestimate the extent and diversity of aSyn pathology. Multiple well-characterised antibodies are crucial to uncover disease-specific features and develop biomarkers for synucleinopathies (Baba et al., 1998; Covell et al., 2017; Dhillon et al., 2017; Duda et al., 2000b; Giasson et al., 2000b). If successfully deciphered, these features could be exploited to develop novel approaches to identify disease-specific biomarkers for PD and other synucleinopathies. They will also improve our understanding of the evolution of aSyn pathology and disease progression (Jellinger, 2009; Milber et al., 2012; Parkkinen et al., 2011, 2008; Steiner et al., 2018).

We sought to develop an antibody toolbox that would capture a range of soluble, aggregated and post-translationally modified forms of aSyn. First, a combination of peptides and wild-type (WT) or site-specifically modified aSyn proteins were designed and used as immunogens to identify antibodies targeting different epitopes and modifications along the sequence of aSyn. The newly generated antibodies were complemented with a pre-existing antibody selection, and these were systematically characterised with regards to their a) epitopes and species reactivity; b) sensitivities to aSyn PTMs neighbouring their epitopes; c) reactivities to other amyloidogenic proteins; and d) conformational selectivity using a library of recombinant aSyn monomers, oligomers and fibrils (Figure 1D). We validated their specificity to aSyn by employing aSyn knockout (KO) mouse neurons and brain tissues. The antibodies were then titrated for use on post-mortem synucleinopathy tissues. We selected an antibody subset that would cover the N-terminus, non-amyloid component (NAC) and C-terminus regions of aSyn as well as the post-translationally modified aSyn forms to profile the pathological lesions in sporadic and familial LBDs. This approach revealed distinct and heterogeneously modified aSyn pathologies rich in pS129, but also in N-terminal nitration at Tyrosine 39 (Y39) and in N- and C-terminal tyrosine phosphorylations scattered both to neurons and glia. To the best of our knowledge, this is the first study to employ antibodies targeting the key disease-associated PTMs to assess aSyn brain pathology in the same set of LBD cases. Finally, we assessed the distribution of differentially modified aSyn aggregates in seeding-based cellular and animal models, and showed that aSyn may become hyperphosphorylated during the aggregation and inclusion maturation processes. These findings emphasise the importance of comprehensive toolsets to uncover the diversity of aSyn species in human tissues and model organisms, advancing our understanding of aSyn pathology, neurodegeneration, and disease progression.

## RESULTS

### Design, development and generation of aSyn monoclonal antibodies

To develop antibodies that capture the diverse biochemical and structural variations of aSyn, we employed a range of human aSyn proteoforms as antigens. These included large peptide fragments and proteins spanning the N-terminal, NAC and C-terminal regions and/or containing pathology-associated PTMs of aSyn (Supplementary Table 1). To ensure that all the disease-associated C-terminal truncations can be targeted, we included antigen sequences encompassing aSyn residues 108-120, 113-127 or 108-140. To increase the likelihood of generating antibodies with epitopes in the N-terminal and/or the NAC region of the protein, we used aSyn 1-20 peptide as well as human aSyn full-length (1-140) and aSyn 1-119 recombinant proteins as immunogens. For the generation of aSyn pS129 antibodies, we initiated two programmes – one with human aSyn 124-135 peptide phosphorylated at S129 (aSyn 124-135 pS129), and another with human aSyn 120-135 peptide doubly phosphorylated at Y125 and S129 (aSyn 120-135 pY125/pS129). Our goal was to develop phospho-specific antibodies that would detect aSyn pS129 even in the presence of neighbouring PTMs. Lastly, to produce monoclonal antibodies against this N-terminal PTM, the mice were immunised with aSyn 34-45 peptide nitrated at Y39 (aSyn 34-45 nY39). Following the immunisation of the Bagg’s ALBino colour (BALB/c) mice, test bleeds, hybridoma supernatants and subclones were analysed via enzyme-linked immunosorbent assay (ELISA), dot/slot blot (DB) and Western blot (WB) against a selected library of aSyn proteins (Supplementary Table 2) to determine the strongest and the most specific antibody response. Details of the monoclonal antibody generation process are provided in the Materials and Methods section. A total of 12 aSyn mouse monoclonal antibodies were obtained (Supplementary Table 3), and their purity was validated by sodium dodecyl sulphate polyacrylamide gel electrophoresis (SDS-PAGE)/ Coomassie staining and WB.

### Antibody characterisation using recombinant synuclein proteins and peptides

#### Epitope mapping and sequence specificity

To determine the sequence specificity and epitopes of our antibodies (Figure 2A), we used a library of aSyn recombinant protein standards (Supplementary Table 2). The two antibodies, LASH-EGT403 and 5B10-A12, targeted the N-terminus of aSyn residues 1-5 and 1-10, respectively. All the other novel monoclonal antibodies were mapped to the C-terminal region spanning residues 110-132 and showed staggered coverage of this protein region (Figure 2A-B; Supplementary Table 3). The epitopes of these seven antibodies were mapped to residues 110-115 (2F10-E12), 115-125 (7H10-E12), 120-125 (4E9-G10), 121-125 (1F10-B12), 121-132 (4E9-C12), 123-125 (2C4-B12) and 126-132 (6B2-D12). The results of the epitope mapping of all the novel monoclonal antibodies are summarised in Figure 2B (antibodies marked with *).

**Figure 2:**
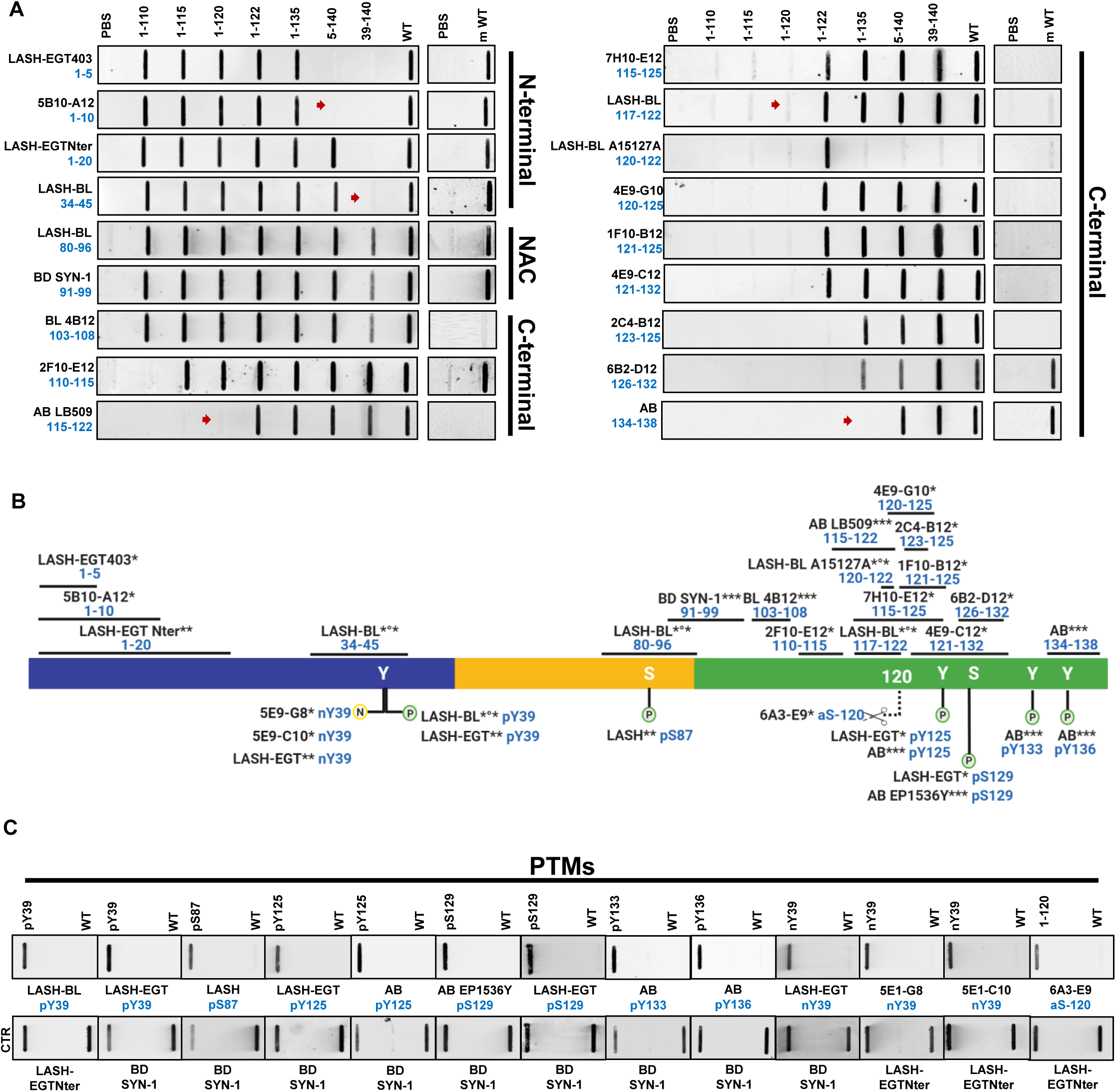
Validation and epitope mapping of aSyn antibodies. **(A)** DB validation of the novel monoclonal, in-house polyclonal and commercially available aSyn antibodies against the N-terminal, NAC and the C-terminal regions of aSyn for epitope mapping, specificity and species reactivity using a selected library of aSyn = recombinant proteins under native conditions. Protein loading control was run via Ponceau S staining. All loaded proteins represent human aSyn forms except for mouse aSyn wild-type (m WT) protein. Red arrows highlight the sensitivities of the antibodies to the presence of neighbouring aSyn PTMs. **(B)** A schematic to represent the novel monoclonal (marked with *), in-house polyclonal (marked with **) and pre-existing commercial aSyn antibodies (marked with ***) included in this study. The commercial antibodies developed jointly with Biolegend are marked with *°*. The epitope information of each antibody is indicated in blue. Schematic created with BioRender.com (agreement no: *JR23G6G5LA*). **(C)** Specificity validation of the aSyn PTM antibodies via DB screening. aSyn = alpha-synuclein; CTR = control; DB = dot/slot blot; FL = full-length; m = mouse; NAC = non-amyloid component; PBS = phosphate buffered saline; PTM = post-translational modification; WT = wild-type

The newly produced monoclonal antibodies alone were not sufficient to cover all regions of the aSyn sequence. Therefore, we incorporated in-house polyclonal aSyn antibodies (Figure 2B, antibodies marked with **), as well as commercially available and frequently used aSyn antibodies (Figure 2B, antibodies marked with ***) within the study. Of these commercial antibodies, LASH-BL 34-45, LASH-BL 80-96, LASH-BL 117-122, LASH-BL A15127A (120-122) and LASH-BL pY39 were developed jointly with Biolegend (Figure 2B, antibodies marked with *°*). The epitopes and sequence specificities of all these antibodies were validated by DB (Figure 2A). We mapped the epitope of the homemade polyclonal LASH-EGTNter antibody to aSyn residues 1-20. LASH-BL A15127A showed a strong preferential positivity to aSyn truncated at residue 122. However, further validation is needed to confirm if this antibody is specific to truncated aSyn (1-122; Figure 2A). For all the other commercial antibodies, the epitopes we identified were similar to those reported in the literature (Figure 2A-B).

In addition to the antibodies against non-modified aSyn, we also generated novel monoclonal antibodies that target specific aSyn PTMs. Three aSyn PTM antibodies were developed, two of which were mapped to aSyn nY39 (5E9-G8 and 5E9-C10), and one to aSyn truncated at residue 120 (6A3-E9) by DB and WB (Figure 2B-C; Supplementary Figure 1A). Amongst the monoclonal antibodies against aSyn nY39, 5E1-G8 was weakly positive to aSyn WT and aSyn nY39 by WB (Supplementary Figure 1A, red arrow) but not by DB (Figure 2C). On the other hand, the other monoclonal antibody 5E1-C10 was positive only for aSyn nY39 both by DB and WB (Figure 2C; Supplementary Figure 1A). The specificity of 6A3-E9 was further validated via supplementary DB, WB and surface plasmon resonance (SPR) analyses (Supplementary Figure 1B-D), all of which showed that it detects human aSyn cleaved at residue 120, but not the full-length protein. We complemented our battery of antibodies against the aSyn PTMs with the previously generated in-house polyclonal antibodies against aSyn nY39 (LASH-EGT nY39), aSyn pY39 (LASH-EGT pY39), aSyn pS87 (LASH pS87), aSyn pY125 (LASH-EGT pY125), aSyn pS129 (LASH-EGT pS129), with the monoclonal aSyn pY39 antibody generated in collaboration with Biolegend (LASH-BL pY39) and with the commercially available aSyn pY125 (AB pY125), pS129 (AB EP1536Y), pY133 (AB pY133) and pY136 (AB pY136) antibodies (Figure 2B). These were validated and were found to be specific to their targeted modifications by DB (Figure 2B-C). Altogether, this selection process provided us with 18 aSyn non-modified and 13 aSyn PTM antibodies to work with. All antibodies used in this study are listed in Supplementary Table 4.

#### Species reactivity

We next assessed the ability of the antibodies to recognise human and mouse aSyn. All N-terminal and NAC-region antibodies, including LASH-EGT403 (1-5), 5B10-A12 (1-10), LASH-EGTNter (1-20), LASH-BL (34-45), LASH-BL (80-96) and BD SYN-1 (91-99) recognised both human- and mouse aSyn (Figure 2A). The large majority of the C-terminal antibodies, on the other hand, detected only human aSyn proteins, with the exception of 2F10-E12 (110-115), 6B2-D12 (126-132) and Abcam (134-138) antibodies that recognised both human and mouse aSyn (Figure 2A). These findings are consistent with the sequence differences between human and mouse aSyn in the C-terminus, particularly in the region covering residues 115-125 (Supplementary Figure 2A).

#### Sensitivity to neighbouring aSyn PTMs

We have recently shown that the aSyn pS129 antibody binding could be affected by the presence of aSyn PTMs neighbouring S129 (Lashuel et al., 2022). Therefore, we investigated the sensitivity of the non-modified aSyn antibodies to the presence of PTMs in close proximity to their epitopes by DB, and the results are summarised in Table 1. A large number of the aSyn C-terminal antibodies (i.e. 8 out of 12) showed sensitivity to the presence of aSyn pY125 and/or truncations within residues 120-125 (Table 1). When aSyn pY125 was present, for instance, antibodies 7H10-E12 (115-125), 4E9-G10 (120-125), 4E9-C12 (121-132) and 2C4-B12 (123-125) failed to produce a strong positive result by DB (Supplementary Figure 2C, red arrows). This finding is in line with the mapped epitopes of these antibodies (Figure 2B), which cover Y125. Therefore, phosphorylation of this residue may interfere with antibody recognition of aSyn. Similarly, the 6B2-D12 (126-132) antibody did not detect aSyn pS129 protein (Supplementary Figure 2C, red arrow). The positive signal revealed by AB LB509 (115-122) and LASH-BL 117-122 were weaker when aSyn was truncated at residue 120 (Figure 2B, red arrows) or when C-terminal residues 120-125 were absent (Supplementary Figure 2C, red arrows). AB 134-138 antibody did not show any positive signal when aSyn was truncated at residue 135 (Figure 2B, red arrow) but was not affected by the presence of phosphorylation at Y136 (Supplementary Figure 2C). We expanded the same analysis to the N-terminal and NAC region antibodies (Table 1). 5B10-A12 (1-10) showed no positivity when aSyn was truncated at residue 5 (Figure 2A, red arrow), which is consistent with the epitope identification of this antibody. The LASH-BL 34-45 antibody showed no sensitivity to the presence of nY39 (Supplementary Figure 2C) but failed to detect aSyn when it was truncated at residue 39 (Figure 2A, red arrow). Detection of aSyn by the other antibodies was not affected by the presence of neighbouring PTMs (Table 2). These results highlight that the antibody sensitivities to the presence of PTMs deserve consideration before aSyn antibodies are selected for prospective experimental studies.

**Table 1:**
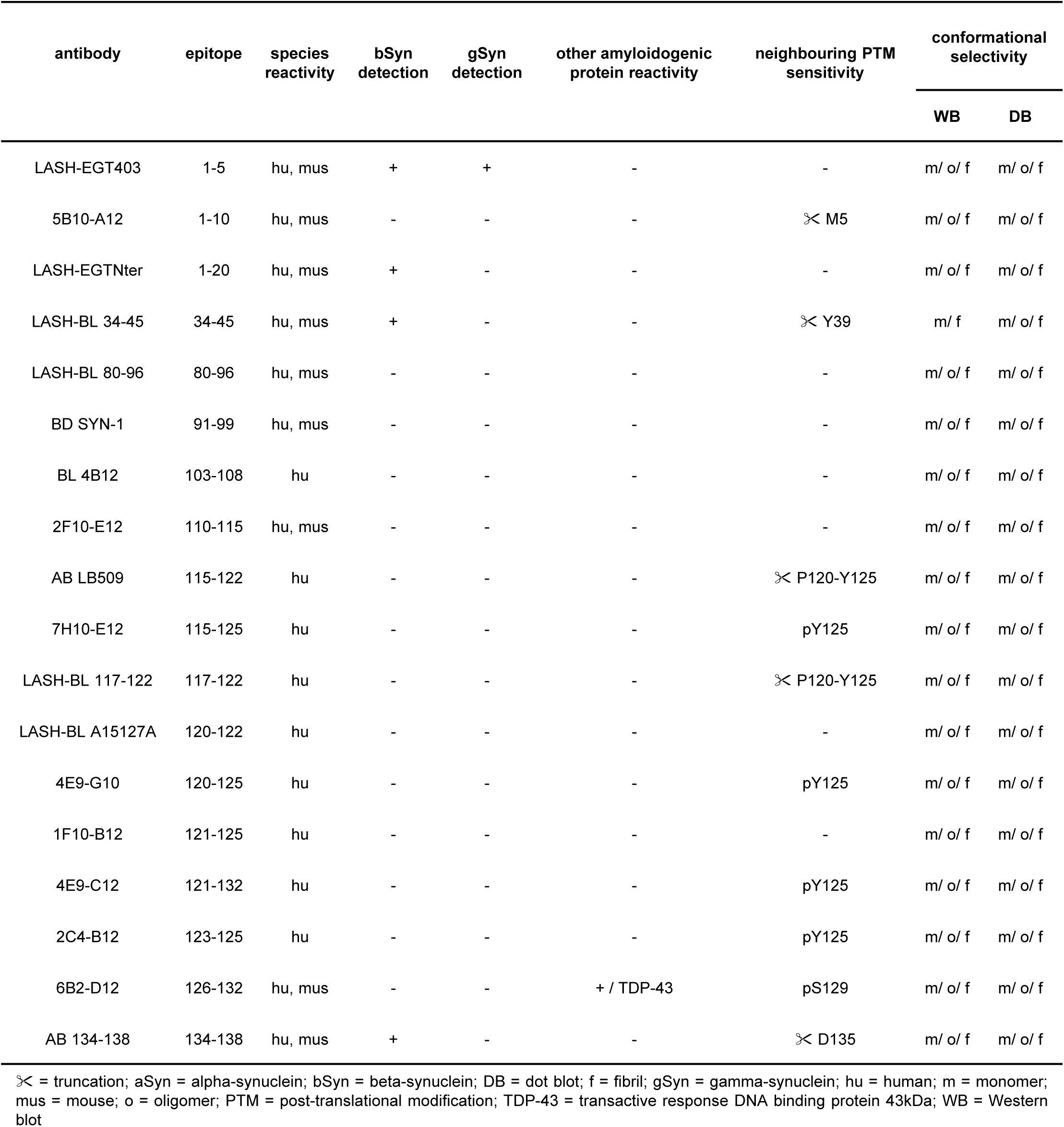
Characterisation of non-modified aSyn antibodies using recombinant synuclein proteins and peptides. ∀ = truncation; aSyn = alpha-synuclein; bSyn = beta-synuclein; DB = dot blot; f = fibril; gSyn = gamma-synuclein; hu = human; m = monomer; mus = mouse; o = oligomer; PTM = post-translational modification; TDP-43 = transactive response DNA binding protein 43kDa; WB = Western blot

**Table 2:** Specificity summary of aSyn antibodies on aSyn KO mouse neuronal culture and brain tissues. + = specific; - = non-specific; aSyn = alpha-synuclein; bSyn = beta-synuclein; hipp = hippocampus; ICC = immunocytochemistry; IHC = immunohistochemistry; kDa = kilodalton; KO = knockout; WB = Western blot

| antibody | epitope | specificity<br>aSyn KO neurons - ICC | specificity<br>aSyn KO neurons – WB |  | specificity<br>aSyn KO tissue – IHC |
| --- | --- | --- | --- | --- | --- |
|  |  |  | 15kDa | 35-180kDa |  |
| LASH-EGT403 | 1-5 | + | + | - | + |
| 5B10-A12 | 1-10 | + | + | - | + |
| LASH-EGTNter | 1-20 | + / bSyn-positive | + | - | + |
| LASH-BL 34-45 | 34-45 | + / bSyn-positive | + | - | + |
| LASH-BL 80-96 | 80-96 | - / hipp aSyn KO only | + | - | + |
| BD SYN-1 | 91-99 | + | + | + | + |
| BL 4B12 | 103-108 | - / mild nuclear | + | - | + |
| 2F10-E12 | 110-115 | + | + | - | + |
| AB LB509 | 115-122 | + | + | - | + |
| 7H10-E12 | 115-125 | - | + | + | + |
| LASH-BL 117-122 | 117-122 | - | + | + | + |
| LASH-BL A15127A | 120-122 | - | + | - | + |
| 4E9-G10 | 120-125 | + | + | - | + |
| 1F10-B12 | 121-125 | - | + | + | + |
| 4E9-C12 | 121-132 | + | + | + | + |
| 2C4-B12 | 123-125 | - | + | + | + |
| 6B2-D12 | 126-132 | + | + | - | + |
| AB 134-138 | 134-138 | + / bSyn-positive | + / bSyn-positive | - | + |
| LASH-BL pY39 | pY39 | + | + | - | + |
| LASH-EGT pY39 | pY39 | + | + | - | + |
| LASH pS87 | pS87 | - / mild nuclear | + | + | - / punctate cytoplasmic |
| LASH-EGT pY125 | pY125 | + | + | + | - / punctate cytoplasmic |
| AB pY125 | pY125 | + | + | + | + |
| AB EP1536Y | pS129 | + | + | + | + |
| LASH-EGT pS129 | pS129 | + | - | - | - / punctate cytoplasmic |
| AB pY133 | pY133 | + | + | + | + |
| AB pY136 | pY136 | + | + | + | + |
| LASH-EGT nY39 | nY39 | + | + | + | + |
| 5E1-G8 | nY39 | - | + | - | - / mild diffuse background |
| 5E1-C10 | nY39 | - | + | - | - / mild diffuse background |
| 6A3-E9 | aSyn-120 | + | + | + | + |

#### Reactivity to synuclein family proteins and other amyloidogenic proteins

The synuclein family of proteins consists of aSyn, beta-synuclein (bSyn) and gamma-synuclein (gSyn; Supplementary Figure 2B) (George, 2001), which share significant sequence homology. We investigated the reactivity of the antibodies targeting non-modified aSyn to bSyn and gSyn. We note that LASH-EGT403 (1-5) was the only antibody to detect aSyn, bSyn and gSyn, therefore acting as a pan-synuclein antibody (Supplementary Figure 2D, blue and green arrows). This is consistent with the fact that the first 4 residues of all the synuclein proteins are identical (Supplementary Figure 2B). The LASH-EGTNter (1-20), LASH-BL (34-45) and the extreme C-terminal antibody AB (134-138) showed reactivity also to bSyn (Supplementary Figure 2D, blue arrows) but not to gSyn. The N-terminal antibody 5B10-A12 (1-10), the NAC-region antibodies and the rest of the C-terminal antibodies did not recognise bSyn or gSyn (Supplementary Figure 2D). These results are in line with the sequence similarities and differences between aSyn, bSyn and gSyn (Supplementary Figure 2B).

We next assessed the specificity of the antibodies against other amyloidogenic proteins including tubulin-associated unit (tau), amyloid-beta (a-beta) and transactive response DNA-binding protein 43kDa (TDP-43). None of the aSyn antibodies revealed any positivity to these amyloidogenic proteins (Supplementary Figure 2D), except for 6B2-D12 (126-132), which was positive for recombinant TDP-43 by DB (Supplementary Figure 2D, red arrow). These results confirmed that 17 out of 18 non-modified aSyn antibodies are specific for recombinant synuclein and do not recognise other amyloidogenic recombinant proteins.

#### Conformational selectivity

aSyn exists in different conformations in healthy and diseased human tissues (Covell et al., 2017; Gai et al., 2003; Kovacs et al., 2012; Roberts et al., 2015; Spillantini et al., 1998a, 1998b). To determine the selectivity of the antibodies towards the different aSyn conformations, we used DB and WB to assess their ability to recognise purified recombinant aSyn WT monomers, oligomers and fibrils (Figure 3A-B; Table 1), which were prepared and characterised as previously described (Kumar et al., 2020b). Electron microscopy (EM) analysis confirmed the aggregation state of aSyn in all three preparations (Figure 3A). All antibodies recognised the three forms of aSyn by DB but showed differential ability to detect SDS-resistant high molecular weight (HMW) oligomers by WB. Namely, LASH-BL 34-45, which detected aSyn monomers, oligomers and fibrils by DB, did not recognise HMW aSyn oligomers under denatured conditions (Figure 3B, red arrow). These results suggest that the vast majority of the selected antibodies are capable of recognising different conformations and aggregation states of aSyn.

**Figure 3:**
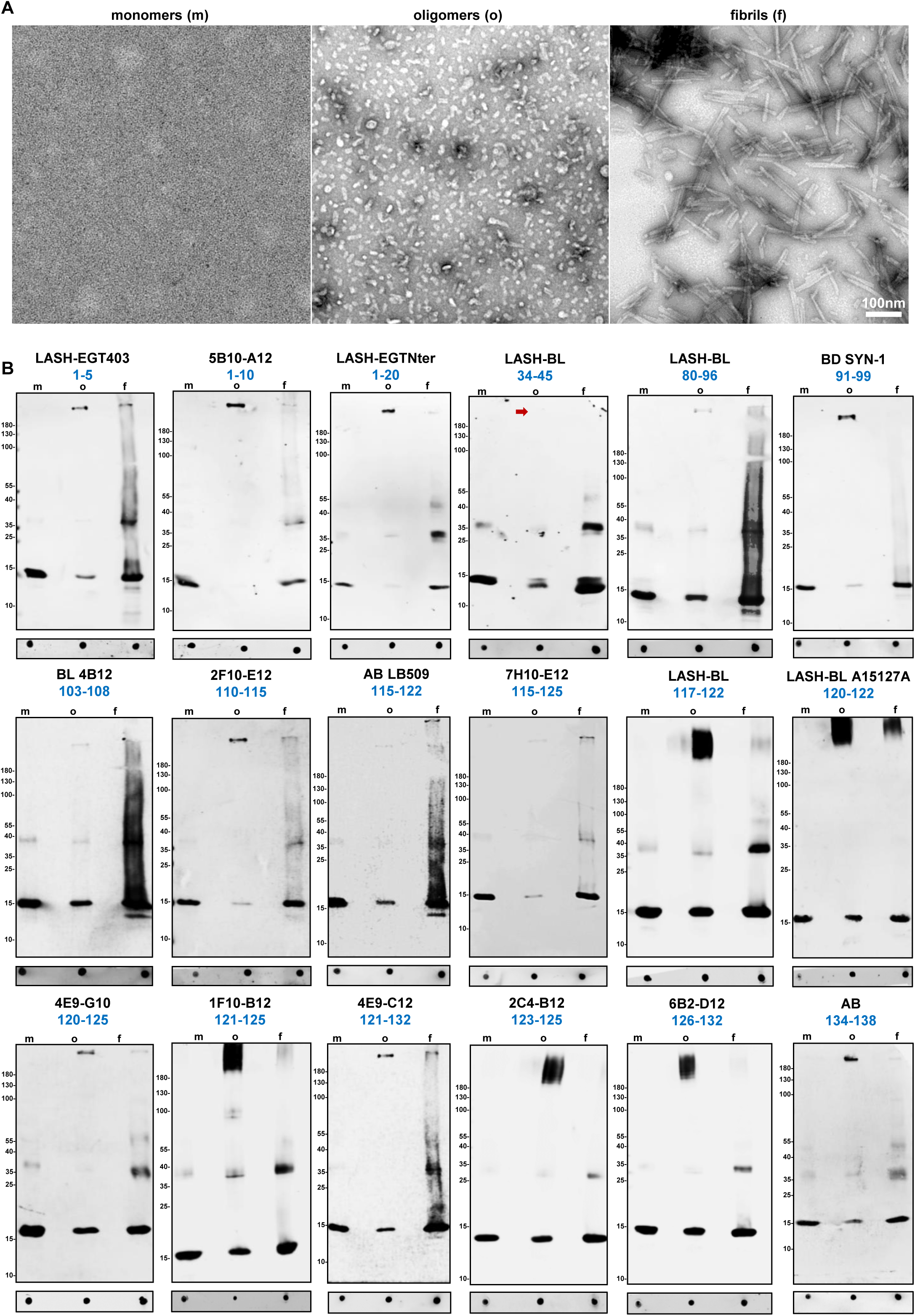
Selectivity of aSyn antibodies over aSyn conformations. **(A)** Representative EM images of aSyn human WT monomers, oligomers and fibrils. **(B)** DB and WB characterisation of 18 aSyn antibodies to determine their conformational selectivity using aSyn human WT recombinant monomers, oligomers and fibrils. aSyn = alpha-synuclein; DB = dot/slot blot; EM = electron microscopy; f = fibrils; m = monomers; o = oligomers; WB = Western blot; WT = wild-type

### Antibody validation using aSyn KO mouse primary neurons and brain tissues

#### aSyn KO mouse primary neurons

To validate the specificity of the antibodies to aSyn species in cells, we assessed their immunoreactivity using primary neurons derived from C57BL/6J-OlaHsd mice (*SNCA*−/−) lacking expression of mouse aSyn. We first determined the minimal working concentration for each of the mouse-reactive antibodies on WT naïve and mouse aSyn pre-formed fibril(PFF)-seeded hippocampal neurons by immunocytochemistry (ICC). This approach allowed us to work on aSyn KO neurons with appropriate antibody concentrations that otherwise permit the detection of endogenous aSyn, exogenously added aSyn fibrils and the newly formed aSyn aggregates in WT neurons, based on a well-characterised neuronal seeding model (Mahul-Mellier et al., 2020, 2018; Volpicelli-Daley et al., 2014). For the antibodies that are human-reactive only, we opted for the recommended dilutions by the suppliers when such information was available, and if not, we aimed at a final antibody concentration of 2-5µg/mL for ICC. We then screened the antibodies using aSyn KO hippocampal and cortical neurons.

By ICC, LASH-EGT403 (1-5), 5B10-A12 (1-10), BD SYN-1 (91-99), 2F10-E12 (110-115), AB LB509 (115-122), 4E9-G10 (120-125), 4E9-C12 (121-132) and 6B2-D12 (126-132) antibodies showed no non-specific background in hippocampal (Supplementary Figure 3A) and cortical (Supplementary Figure 3B) aSyn KO neurons. The LASH-EGTNter (1-20), LASH-BL (34-45) and AB (134-138) antibodies were positive, possibly for bSyn protein, both in hippocampal and cortical aSyn KO neurons (Supplementary Figure 3A-B, blue arrows), an observation consistent with DB results on recombinant bSyn reactivity using these antibodies (Table 1; Supplementary Figure 2D, blue arrows). Collectively, all of these antibodies showed specificity by ICC on aSyn KO hippocampal and cortical neurons. The cytoplasmic and/or nuclear signals detected in hippocampal and cortical aSyn KO neurons with BL 4B12 (103-108), 7H10-E12 (115-125), LASH-BL (117-122), LASH-BL A15127A (120-122), 1F10-B12 (121-125) and 2C4-B12 (123-125), on the other hand, were deemed to be non-specific (Supplementary Figure 3A-B, red arrows). Interestingly, the LASH-BL 80-96 antibody revealed a weak, non-specific cytoplasmic signal in the hippocampal (Supplementary Figure 3A, red arrow) but not in cortical aSyn KO neurons (Supplementary Figure 3B).

Next, we assessed the specificity of the aSyn PTM antibodies in aSyn KO neurons by ICC. The LASH-BL pY39, LASH-EGT pY39, LASH-EGT pY125, AB pY125, AB EP1536Y pS129, LASH-EGT pS129, AB pY133, AB pY136, LASH-EGT nY39 and 6A3-E9 antibodies were negative by ICC (Supplementary Figure 4A-B), suggesting that these antibodies are specific and show no cross-reactivity. In contrast, the monoclonal aSyn nY39 antibodies 5E1-G8 and 5E1-C10 were strongly positive both in the hippocampal and cortical aSyn KO neurons, showing non-specific background. Mild nuclear positivity was observed with LASH pS87 (Supplementary Figure 4A-B, red arrows).

We used a similar approach to investigate the specificity of the aSyn antibodies in neuronal lysates by WB. Sequential extraction was run on aSyn KO hippocampal and cortical neurons, and the soluble and insoluble fractions were profiled using the aSyn antibodies. aSyn mouse or human recombinant standards were used as positive controls. Of the human-reactive non-modified aSyn antibodies, faint non-specific bands were present mainly in the HMW regions of the soluble and/or insoluble fractions of BL 4B12 (103-108), AB LB509 (115-122), LASH-BL A15127A (120-122), 4E9-G10 (120-125) (Supplementary Figure 5A, red arrows), but not in the region where monomeric aSyn migrates, at 15kDa. Non-specific bands were not observed with the human-reactive 7H10-E12 (115-125), LASH-BL (117-122), 1F10-B12 (121-125), 4E9-C12 (121-132) and 2C4-B12 (123-125) antibodies (Supplementary Figure 5A). The human- and mouse-reactive non-modified aSyn antibodies i.e. LASH-EGT403 (1-5), 5B10-A12 (1-10), LASH-EGTNter (1-20), LASH-BL (34-45), LASH-BL (80-96), BD SYN-1 (91-99), 2F10-E12 (110-115), 6B2-D12 (126-132) and AB (134-138), did not detect any non-specific bands at 15kDa in the phosphate buffered saline(PBS)-treated soluble or insoluble fractions (Supplementary Figure 5A), with AB (134-138) possibly detecting bSyn in the PBS-treated soluble neuronal fractions (Supplementary Figure 5A, blue arrows). On the other hand, LASH-EGT403, 5B10-A12, LASH-EGTNter, LASH-BL 34-45, LASH-BL 80-96, 2F10-E12, 6B2-D12 and AB 134-138 showed non-specific bands in the HMW regions in the soluble and/or insoluble fractions (Supplementary Figure 5A, red arrows). Interestingly, most of the aSyn PTM antibodies did not reveal any non-specificity, except for LASH-BL pY39, LASH-EGT pY39, 5E1-G8 nY39 and 5E1-C10 nY39 that detected a few non-specific bands between 35-180kDa, and LASH-EGT pS129 which detected both HMW bands and bands around 15kDa in the insoluble fractions (Supplementary Figure 5B, red arrows). We summarised these results in Table 2.

#### aSyn KO mouse brain tissues

Next, we validated the specificity of the aSyn antibodies on aSyn KO mouse amygdala tissue, a brain region previously shown to be particularly affected by aSyn pathology (Burtscher et al., 2019). By immunofluorescence (IF), there was no background with any of the antibodies against non-modified aSyn (Supplementary Figure 6A). With the aSyn PTM antibodies, on the other hand, we observed non-specific punctate cytoplasmic positivity with LASH pS87, LASH-EGT pY125, LASH-EGT pS129 and mild diffuse background with 5E1-G8 and 5E9-C10 nY39 antibodies (Supplementary Figure 6B, red arrows). A summary of the findings on aSyn KO brain tissue for each antibody is presented in Table 2.

### Antibody validation and application in human post-mortem brain tissues

Following the validation of the antibodies using recombinant proteins, aSyn KO mouse primary neurons and brain samples, we applied these tools on human post-mortem formalin-fixed paraffin-embedded (FFPE) brain tissues. The antibodies were titrated for immunohistochemistry (IHC) by comparing different pre-treatment (i.e. epitope retrieval) conditions and antibody dilutions. All of the non-modified aSyn antibodies (18) revealed moderate to extensive staining of aSyn pathology in the PD cingulate cortex (Figure 4A) except for the LASH-EGT403 (1-5) and 6B2-D12 (126-132) antibodies, which were rarely immunoreactive to LBs and did not detect LNs.

**Figure 4:**
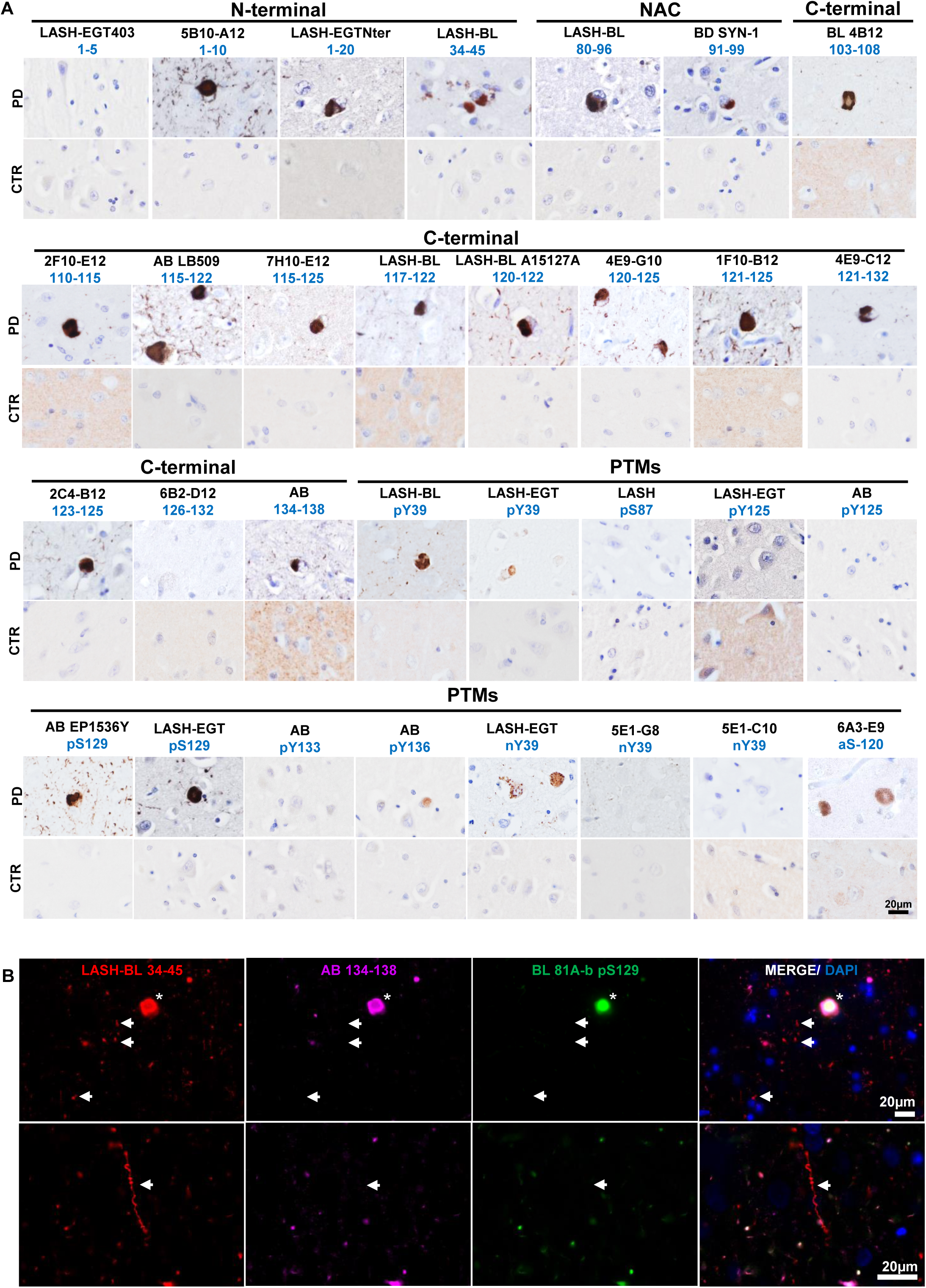
Application and validation of the aSyn antibodies on PD tissue. **(A)** The in-house monoclonal, polyclonal and commercial aSyn antibodies were optimised for IHC on the PD cingulate cortex. Non-specific staining was not observed in age-matched healthy controls. Representative images taken from the cortical deep grey matter (layers V–VI) of PD1, CTR1 and CTR2. **(B)** Triple labelling of PD cingulate cortex by IF using an aSyn N-terminal (LASH-BL 34-45), a C-terminal (AB 134-138) and a pS129 (BL 81A-biotin) antibody. LBs are marked with asterisks, and LNs with arrows. Representative images from PD1 cingulate cortex are taken using Leica DM5500 B upright microscope at 20x magnification. aSyn = alpha-synuclein; CTR = control; DAPI = 4’,6-diamidino-2-phenylindole; IF = immunofluorescence; IHC = immunohistochemistry; LB = Lewy body; LN = Lewy neurite; NAC = non-amyloid component; PD = Parkinson’s disease; PTM = post-translational modification

Amongst the three PTM antibodies against aSyn nY39, only the LASH-EGT nY39 polyclonal antibody positively detected cortical LBs and LNs. The 5E1-G8 and 5E1-C10 nY39 monoclonal antibodies, on the other hand, were negative on PD tissue by IHC. With regards to aSyn phosphorylation, some LBs but also thin neurites were positive for aSyn pY39, detected both by the monoclonal LASH-BL pY39 and by the polyclonal LASH-EGT pY39 antibodies. Likewise, extensive LB and LN pathology was revealed both by the commercial AB EP1536Y and the homemade LASH-EGT pS129 antibodies. On the contrary, the LASH pS87 and LASH-EGT pY125 antibodies showed little to no reactivity in the PD cingulate cortex. Whilst we could not detect any positivity in PD cingulate cortex with AB pY125 and AB pY133, we found these antibodies to reveal LBs and neuritic pathology in DLB entorhinal cortex (Altay et al., 2022). This suggests that these antibodies specifically reveal the C-terminal tyrosine phosphorylations on post-mortem human tissues when this modification is present. The truncation-specific antibody 6A3-E9 (aSyn-120) detected exclusively the LBs without producing signal in the neurites (Figure 4A).

Interestingly, triple immunolabelling using aSyn N-terminal, C-terminal and PTM antibodies revealed that the cortical LBs were equally detected by all antibodies (Figure 4B; Supplementary Figure 7, asterisks). Yet, some of the LNs and neuropil dots were selectively revealed by the LASH-BL 34-45 antibody, and not by the AB 134-138 or the 81A pS129 antibodies (Figure 4B; Supplementary Figure 7 arrows), suggesting that a portion of neuritic pathology is non-phosphorylated and may be cleaved in the extreme C-terminus.

### Panel selection of aSyn antibodies reveals the biochemical and morphological diversity of human aSyn pathology

In human tissues, aSyn displays heterogeneous conformations and proteoforms (Anderson et al., 2006; John E Duda et al., 2000; John E. Duda et al., 2000; Fujiwara et al., 2002; Giasson et al., 2000b; Kuusisto et al., 2003). To further investigate aSyn pathology in various types of LBDs, we carefully selected a subset of highly specific and effective antibodies that offers extensive coverage of the aSyn sequence and its post-translational modifications. We included two antibodies against the N-terminus (LASH-EGTNter 1-20 and LASH-BL 34-45), two antibodies against the NAC region (LASH-BL 80-96 and BD SYN-1 91-99), and two antibodies against the C-terminus (i.e. 2F10-E12 110-115 and AB 134-138) of aSyn. Two antibodies were incorporated to cover the aSyn serine phosphorylations: AB EP1536Y pS129, as this antibody has been the most specific to aSyn pS129 species in our hands and in the literature (Delic et al., 2018; Lashuel et al., 2022), and LASH pS87, which was recently shown to detect LBs in PD and GCIs in MSA (Sonustun et al., 2022). For the N-terminal tyrosine phosphorylation modification, we opted for the monoclonal LASH-BL pY39 antibody. For the C-terminal phosphorylation at Y125, we selected the polyclonal aSyn pY125 antibody from Abcam as it did not show non-specific background signal in aSyn KO mouse brain tissues (Supplementary Figure 6B). To cover all possible C-terminal tyrosine phosphorylation modifications, we also included the polyclonal aSyn pY133 and pY136 antibodies from Abcam. To the best of our knowledge, these two C-terminal tyrosine phosphorylations have not been explored in post-mortem tissues, with the exception of one recent study (Sano et al., 2021). Lastly, we added 6A3-E9 against aSyn 1-120 and LASH-EGT nY39 to our selection. The final antibody subset (14) chosen for further human tissue screening is shown in Figure 5A.

**Figure 5:**
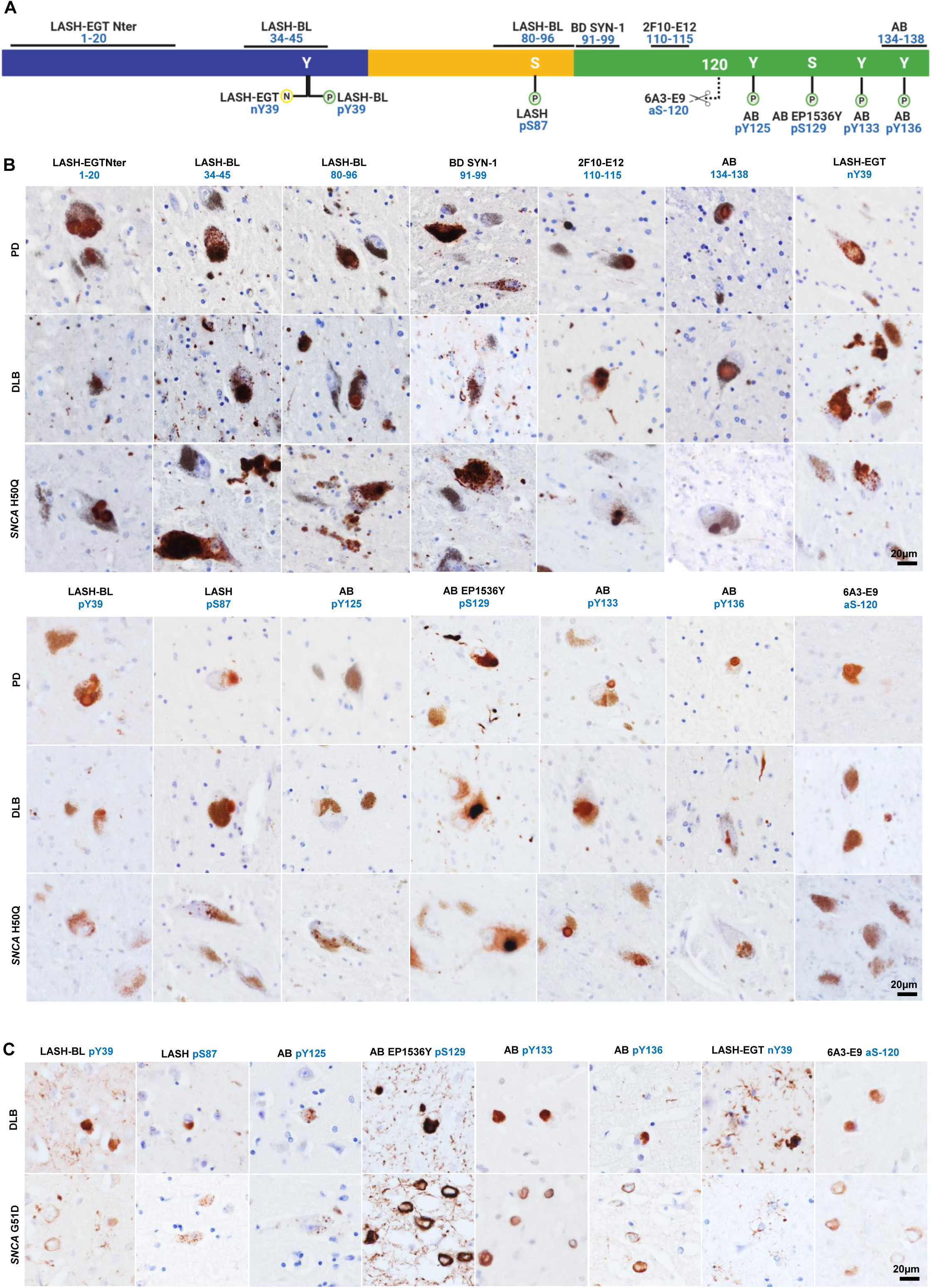
A selected panel of aSyn antibodies reveal the broad diversity of human pathology in the SN of LBDs. **(A)** An outline to show the epitopes of the aSyn antibody selection used for IHC studies on LBD tissues. Schematic created with BioRender.com (agreement no: *NU23G6E7KK*). **(B)** Representative images from the substantia nigra of sporadic (PD, DLB) and familial (*SNCA* H50Q) LBDs screened with the selection of aSyn non-modified and PTM antibodies. Images taken from the SN of PD2, PD3, DLB1 and *SNCA* H50Q1. **(C)** Representative images from the cingulate cortex of sporadic (DLB) and familial (*SNCA* G51D) LBDs screened with the selected aSyn PTM antibodies. Images taken from the cortical deep grey matter (layers V–VI) of DLB1 and *SNCA* G51D1. aSyn = alpha-synuclein; DLB = dementia with Lewy bodies; IHC = immunohistochemistry; LBD = Lewy body disease; SN = substantia nigra; PD = Parkinson’s disease; PTM = post-translational modification

We validated the specificity of these 14 antibodies to aSyn species directly in human brain tissue using post-mortem brain samples with neurodegenerative diseases that are not classified as synucleinopathies. The frontal cortices of progressive supranuclear palsy (PSP) and corticobasal degeneration (CBD), and the hippocampi and entorhinal cortices of Alzheimer’s disease (AD), Pick’s disease (PiD) and frontotemporal lobar degeneration of TDP-43 type C (FTLD-TDP/C) cases were immunostained. None of the non-modified aSyn antibodies showed any immunoreactivity (Supplementary Figure 8). With regards to the aSyn PTM antibodies, LASH-EGT nY39, LASH-BL pY39, AB pY133 and 6A3-E9 aSyn-120 were negative in all cases stained (Supplementary Figure 8). LASH pS87 and AB pY125 detected sparse granular cytoplasmic aSyn, and AB EP1536Y (pS129) some neuritic aSyn in AD entorhinal cortex (Supplementary Figure 8, arrows). In addition, LASH pS87 revealed rare cytoplasmic positivity in PiD and AB pY136 in FTLD-TDP/C hippocampi (Supplementary Figure 8, arrows). Given that aSyn accumulation within the inclusions has been reported in AD (Arai et al., 2001), and PiD (Mori et al., 2002) post-mortem brains, we believe that these positive structures may represent genuine aSyn species.

The selected aSyn antibodies were then used to immunohistochemically analyse the substantia nigra of sporadic (PD n=6, PDD n=2, DLB n=1) and familial (*SNCA* G51D n=3, *SNCA* H50Q n=1, *SNCA* duplication n=1) LBDs, and the results summarised in Table 3. The LBs and LNs were uniformly detected by all aSyn non-modified and PTM antibodies except for AB pY125 and 6A3-E9 (aSyn-120) (Figure 5B; Supplementary Figure 9A; Table 3). The neuronal diffuse cytoplasmic aSyn with or without an aggregate were likewise revealed by all antibodies with the exception of LASH-BL pY39 and AB pY136. The neuronal punctate cytoplasmic aSyn species were revealed only by a subgroup of the non-modified aSyn antibodies, namely LASH-BL 34-45, LASH-BL 80-96 and BD SYN-1 (91-99). Similarly, the astroglial accumulations were exclusively revealed by these three non-modified aSyn antibodies and the LASH-BL pY39 and the LASH-EGT nY39 antibodies, and not by any other antibodies included in the study (Figure 5B; Supplementary Figure 9A; Table 3). Interestingly, in the cingulate cortex of these LBDs, we observed a similar phenomenon where LASH-BL pY39 and LASH-EGT nY39 antibodies picked up star-shaped glial structures (Figure 5C; Supplementary Figure 9B). The presence of coiled body-like oligodendroglial accumulations of aSyn in the substantia nigra has been reported previously (Wakabayashi et al., 2000). These oligodendroglial species were revealed best by 2F10-E12 (110-115) (Table 3). The extracellular aSyn species were detected by all aSyn non-modified and PTM antibodies except for AB pY125, and were particularly enhanced with truncation at residue 120 as well as nitration at Y39 (Figure 5B; Supplementary Figure 9A; Table 3).

**Table 3:** A summary of the pathology detection patterns of aSyn non-modified and PTM antibodies on LBD substantia nigra, including the morphological visualisation of these pathological aSyn accumulations. - absent; + mild; ++ moderate; +++ frequent; aSyn = alpha-synuclein; LB = Lewy body; LBD = Lewy body disease; LN = Lewy neurite; PTM = post-translational modification

| antibody | NEURONAL |  |  |  | GLIAL |  | EXTRACELLULAR |
| --- | --- | --- | --- | --- | --- | --- | --- |
|  | LBs | diffuse cytoplasmic with an aggregate | punctate cytoplasmic | LNs & neuropil dots | astroglial | coiled body-like oligodendroglial |  |
| LASH-EGTNter 1-20 | ++ | ++ | - | ++ | - | + | + |
| LASH-BL 34-45 | +++ | + | +++ | +++ | +++ | + | + |
| LASH-BL 80-96 | ++ | + | ++ | ++ | ++ | + | + |
| BD SYN-1 91-99 | ++ | + | ++ | ++ | ++ | + | + |
| 2F10-E12 110-115 | +++ | ++ | - | +++ | - | ++ | + |
| AB 134-138 | ++ | ++ | - | ++ | - | - | + |
| LASH-BL pY39 | + | - | -/+ | + | + | + | + |
| LASH pS87 | + | + | -/+ | + | - | - | + |
| AB pY125 | - | + | -/+ | - | - | - | - |
| AB EP1536Y pS129 | +++ | ++ | - | +++ | - | + | + |
| AB pY133 | + | + | - | + | - | - | + |
| AB pY136 | ++ | - | - | ++ | - | - | + |
| LASH-EGT nY39 | + | + | ++ | + | ++ | - | ++ |
| 6A3-E9 aSyn-120 | - | + | - | - | - | - | ++ |

Both in the substantia nigra and the cingulate cortex, the most abundant aSyn PTM was aSyn pS129, and the AB EP1536Y antibody against this modification labelled LBs, diffuse neuronal cytoplasmic aSyn, neurites and neuropil dots extensively (Figure 5B-C; Supplementary Figure 9A-B). In contrast, LASH pS87 together with AB pY125 revealed only sparse structures across the LBDs. LASH pS87 antibody unveiled very rare puncta in the neuronal cytoplasm, dystrophic neurites and diffuse neuronal cytoplasmic accumulations in the substantia nigra (Figure 5B; Supplementary Figure 9A) but no glial accumulations or thin threads (Figure 5B-C; Supplementary Figure 9A-B). AB pY125 labelled only very rare punctate cytoplasmic aSyn structures and rare diffuse cytoplasmic inclusions in the substantia nigra, without staining any neurites or LBs in either of the two regions examined (Figure 5B-C; Supplementary Figure 9A-B). The other C-terminal tyrosine phosphorylation antibody AB pY133 detected some neurons containing multiple LBs, diffuse neuronal cytoplasmic inclusions and rare thin threads, whereas AB pY136 moderately picked up LBs, dystrophic neurites, neuropil dots and thin threads. The truncation-specific 6A3-E9 antibody (aSyn-120), on the contrary, stained diffuse neuronal cytoplasmic and extracellular aSyn structures, but no neurites. Together, our research revealed a wide spectrum of abnormal aSyn accumulations in both neurons and glial cells within LBD brains. We found that specific modifications of aSyn vary depending on the cell type, and we identified various combinations of modified aSyn species that have not been thoroughly documented before. For instance, we discovered the presence of aSyn nY39, pY133, and pY136, which have not been previously described in a comprehensive manner.

### Biochemical and morphological diversity of aSyn aggregates in cellular and animal seeding models

Seeding-based cellular and animal models using aSyn PFFs have emerged as the most common tools to investigate mechanisms and pathways of aSyn pathology formation (Kumar et al., 2021; Luk et al., 2012, 2009; Mahul-Mellier et al., 2020, 2018; Volpicelli-Daley et al., 2014, 2011). To characterise the diversity of aSyn species in these PFF models, we investigated the ability of our antibodies to detect exogenously added PFFs in aSyn KO hippocampal and cortical neurons by ICC and WB (Supplementary Figure 3-5) 14 hours after treatment. Our findings are summarised in Table 4. All of the 9 non-modified and mouse aSyn-reactive antibodies i.e. the N-terminal LASH-EGT403 (1-5), 5B10-A12 (1-10), LASH-EGTNter (1-20) and LASH-BL 34-45; the NAC-region LASH-BL 80-96, BD SYN-1 (91-99); and the C-terminal 2F10-E12 (110-115), 6B2-D12 (126-132) and AB 134-138 antibodies detected exogenous fibrils in the PFF-treated hippocampal (Supplementary Figure 3A) and cortical (Supplementary Figure 3B) aSyn KO neurons by ICC. By WB, bands specific to exogenously added fibrils were revealed between 10kDa-35kDa by these antibodies (Supplementary Figure 5A, green arrows) except for the 6B2-D12 (126-132) antibody, which failed to detect any bands in the PFF-added neuronal fractions. With the aSyn PTM antibodies, no positivity was detected in the PFF-treated hippocampal or cortical aSyn KO neurons using the aSyn pY39, pS129, pY133 or pY136 antibodies by ICC (Supplementary Figure 4A-B) or by WB (Supplementary Figure 5B). Similarly, no positivity was revealed with the LASH-EGT nY39 antibody in the hippocampal or cortical aSyn KO neurons by ICC (Supplementary Figure 4A-B) or WB (Supplementary Figure 5B). On the contrary, the monoclonal nY39 antibodies 5E1-G8 and 5E1-C10 were positive both in the PFF-treated and control (PBS-treated) aSyn KO neurons by ICC, signals which we presumed to be non-specific (Supplementary Figure 4A-B, red arrows). Intriguingly, 6A3-E9 antibody against aSyn-120 showed no background in PBS-treated control neurons but was positive in PFF-treated hippocampal and cortical aSyn KO neurons by ICC (Supplementary Figure 4A-B, green arrows). Therefore, we cannot rule out the possibility that this antibody may be reactive to aSyn full-length mouse fibrils. By WB, on the other hand, no bands were revealed in the PFF-treated aSyn KO hippocampal and cortical soluble and insoluble fractions (Supplementary Figure 5B). Collectively, these results confirm the specificity of our antibodies, and suggest that the internalised PFFs do not undergo any type of modifications except for N- and C-terminal cleavages, in the absence of seeding.

**Table 4:** A summary of the mouse aSyn-reactive antibody detection patterns of exogenous PFFs in aSyn KO neurons, newly formed aggregates in PFF-seeded WT neurons and mouse brain tissues. *aSyn pS129 revealed by BL 81A mouse monoclonal and AB MJF-R13 rabbit monoclonal antibodies. aSyn = alpha-synuclein; ICC = immunocytochemistry; KO = knockout; PFF = pre-formed fibril; WB = Western blot; WT = wild-type

| antibody | epitope | aSyn KO neurons |  | PFF seeding model:<br>WT primary neurons |  | PFF seeding model:<br>WT mouse brain tissue |  |
| --- | --- | --- | --- | --- | --- | --- | --- |
|  |  | PFF detection<br>ICC | PFF detection<br>WB | positivity | aSyn pS129*<br>overlap | positivity | aSyn pS129*<br>overlap |
| LASH-EGT403 | 1-5 | + | + | + | - | - | - |
| 5B10-A12 | 1-10 | + | + | + | + | + | + |
| LASH-EGTNter | 1-20 | + | + | + | + | + | + |
| LASH-BL 34-45 | 34-45 | + | + | + | + | + | - |
| LASH-BL 80-96 | 80-96 | + | + | + | - | + | - |
| BD SYN-1 | 91-99 | + | + | + | - | + | - |
| 2F10-E12 | 110-115 | + | + | + | - | + | + |
| 6B2-D12 | 126-132 | + | - | + | - | + | - |
| AB 134-138 | 134-138 | + | + | + | + | + | + |
| LASH-EGT pY39 | pY39 | - | - | + | - | + | - |
| AB EP1536Y | pS129 | - | - | + | + | + | + |
| LASH-EGT pS129 | pS129 | - | - | + | + | + | + |
| AB pY133 | pY133 | - | - | + | - | - | - |
| AB pY136 | pY136 | - | - | + | - | - | - |
| LASH-EGT nY39 | nY39 | - | - | - | - | - | - |
| 5E1-G8 | nY39 | +(non-specific) | - | +(non-specific) | - | - | - |
| 5E1-C10 | nY39 | +(non-specific) | - | +(non-specific) | - | - | - |

Finally, the mouse aSyn-reactive antibodies (Table 1; Table 4) were used to characterise the PTM profile of newly formed aSyn aggregates in the neuronal seeding model (Mahul-Mellier et al., 2020, 2020; Volpicelli-Daley et al., 2014, 2011), and in the PFF-injected *in vivo* model (Burtscher et al., 2019; Luk et al., 2012; Masuda-Suzukake et al., 2013) of aSyn. Whilst all the non-modified aSyn antibodies detected the endogenous aSyn in PFF-treated or control (PBS-treated) WT hippocampal neurons by ICC, the 5B10-A12 (1-10), LASH-EGTNter (1-20), LASH-BL 34-45 and AB 134-138 antibodies showed almost complete overlap with the aSyn pS129-positive inclusions in the PFF-treated neurons (Table 4; Figure 6A, arrows), suggesting that these could be useful and alternative tools to aSyn pS129 for monitoring the aSyn aggregation in cell culture, especially if cross-reactivity of the pS129 antibodies is a confounding factor. A similar pattern of overlap with the aSyn-positive inclusions, specifically for 5B10-A12 (1-10), LASH-EGTNter (1-20), 2F10-E12 (110-115) and AB 134-138 antibodies (Table 4; Figure 6C, arrows), was seen in the amygdala of WT mice that had been injected with PFFs in the striatum. The amygdala is particularly prone to develop early and substantial aSyn pathology in this model (Burtscher et al., 2019). We speculate that this staining pattern may be due to the preferential exposure of epitopes in the extreme N- and C-terminal aSyn, whereas the hydrophobic NAC region is less accessible and buried in the core of the newly formed aggregates.

**Figure 6:**
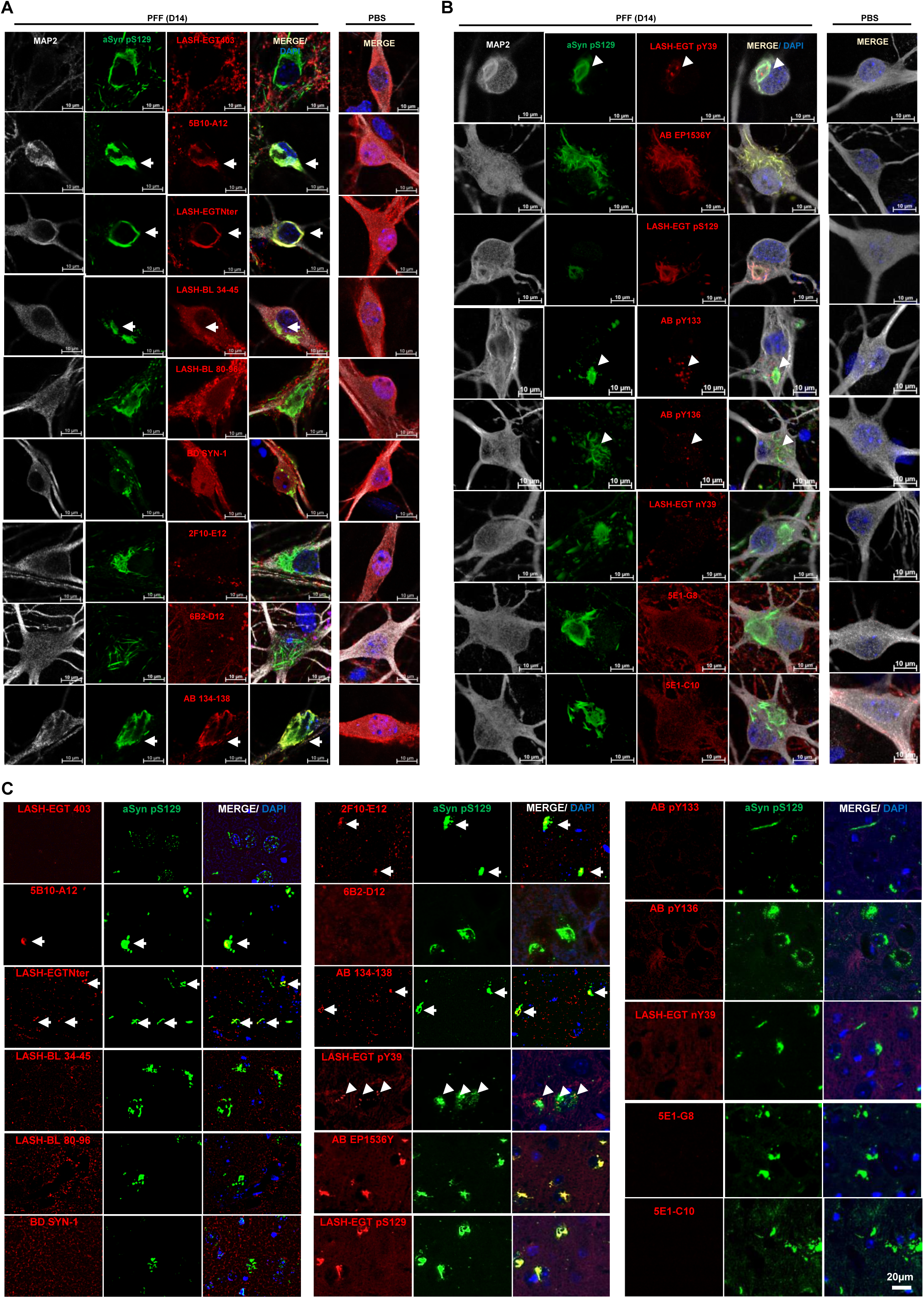
Application of the aSyn antibodies to the cellular and animal seeding models to profile the newly formed aSyn aggregates. WT hippocampal neurons were seeded with PFFs for 14 days, and the newly formed aggregates monitored by ICC using the mouse-reactive **(A)** non-modified aSyn and **(B)** aSyn PTM antibodies in parallel to aSyn pS129 antibodies BL 81A or AB MJF-R13. **(C)** The same type of screening was run in PFF-injected mouse amygdala tissues by IHC. The non-modified aSyn antibody signals overlapping with the aSyn pS129-positive aggregates are marked with an arrow. The punctate positivity shown by aSyn pY39, pY133 and pY136 antibodies in close proximity to aSyn pS129-positive aggregates are shown by arrowheads. Note the non-specific diffuse positivity revealed by the two monoclonal nY39 antibodies 5E1-G8 and 5E1-C10 in the WT hippocampal neurons are also revealed in the aSyn KO neurons using these two antibodies (Supplementary Figure 4A-B). aSyn = alpha-synuclein; DAPI = 4’,6-diamidino-2-phenylindole; ICC = immunocytochemistry; IHC = immunohistochemistry; KO = knockout; MAP2 = microtubule-associated protein 2; PBS = phosphate buffered saline; PFF = pre-formed fibril; PTM = post-translational modification; WT = wild-type

With the aSyn nY39 antibodies, we did not see any positivity in the neurons or in mouse tissue (Table 4; Figure 6B-C), except for diffuse cytoplasmic staining with the 5E1-G8 and 5E1-C10, which we also observed in aSyn KO neurons and therefore deemed non-specific (Supplementary Figure 4A-B). With the N-terminal tyrosine phosphorylation at Y39, on the other hand, both in the PFF-treated neurons and in the PFF-injected mouse amygdala, we detected punctate structures in close proximity to, and partially overlapping with, the aSyn pS129-positive inclusions (Table 4; Figure 6B-C, arrowheads). We noted similar punctate positivity also with aSyn pY133 and pY136 antibodies in the neuronal seeding model (Table 4; Figure 6B, arrowheads), which again partially overlapped with the aSyn pS129-positive accumulations. Altogether, these data suggest that aSyn may become hyperphosphorylated during the aggregation and inclusion maturation processes. Further research is needed to decipher the mechanism of formation and fate of these puncta structures. In this study, we have demonstrated the effectiveness of the antibodies we introduced in examining how PTMs dominate aSyn seeding and inclusion formation. Additionally, our antibodies enable us to capture the diverse structural and chemical characteristics of aSyn aggregates in both cellular and *in vivo* models of aSyn seeding.

## DISCUSSION

Previous studies focusing on the development and application of multiple antibodies have been instrumental in the understanding of pathological heterogeneity of aSyn in LBDs and MSA (Almandoz-Gil et al., 2017; Covell et al., 2017; Croisier et al., 2006; Dhillon et al., 2017; Duda et al., 2002, 2000b; Fagerqvist et al., 2013; Galvin et al., 2000; Giasson et al., 2000a, 2000b; Henderson et al., 2020; Vaikath et al., 2015; Waxman et al., 2008). In this study we developed, characterised and validated 12 novel aSyn antibodies targeting the entire sequence and several disease-associated PTMs of aSyn. These were complemented with pre-existing antibodies to guarantee a more comprehensive coverage of the aSyn biochemical and structural diversity. In total, 31 antibodies were thoroughly characterised in a single study using a stringent validation pipeline (Figure 1D). We selected 14 of these antibodies to profile aSyn pathology on the same set of LBD cases (Figure 5A; Table 3). All of the antibodies targeting non-modified aSyn sequences efficiently detected LBs, but we observed considerable differences, particularly in the revelation of astroglial aSyn accumulations, punctate neuronal cytoplasmic inclusions and a subset of LNs and neuropil dots in the substantia nigra and cingulate cortex of LBDs. This finding suggests differential capacities of the individual antibodies to uncover specific aspects of aSyn pathology with important implications for disease staging and characterisation of disease subtypes.

LBs are mature inclusions consisting of a diverse range of aSyn species, including full-length, N-and C-terminally truncated, and other modified forms with different conformations (Bhattacharjee et al., 2019; Moors et al., 2021; Prasad et al., 2012). Consequently, aSyn antibodies targeting different epitopes throughout its sequence are more likely to detect LBs. However, specific forms or conformations of aSyn in other types of accumulations may be challenging to detect using single antibodies. These include less mature and poorly understood manifestations of aSyn pathology, such as punctate neuronal inclusions, astroglial aSyn accumulations, and a subset of LNs, potentially composed of uniform aSyn proteoforms. By employing an expanded toolbox, we can now systematically analyse aSyn species at the single-cell level using ICC/IHC. Through this approach, we have identified antibodies capable of detecting specific aSyn species and inclusions in different cell types. This advancement opens doors for future investigations into the factors influencing aSyn pathology formation, such as sequence, cell type and cellular environment determinants, as well as allowing the exploration of various cellular and molecular processes associated with neurodegeneration.

Our work allowed for the investigation of the most prominent disease-associated phosphorylation, nitration and truncation PTMs of aSyn at differing densities on the same set of LBD cases. We confirm that aSyn pS129 is the most abundant disease-related modification of aSyn in the substantia nigra, an observation consistent with publications reporting aSyn pS129 enrichment in diseased brains by mass spectrometry analyses (Anderson et al., 2006; Fujiwara et al., 2002). Uniquely, we report here that LB disorders are also rich in N-terminal nitration at Y39. Nitrated aSyn in post-mortem human studies has been previously reported (Duda et al., 2000) but the site-specific N-terminal nitration at Y39 was not investigated due to the lack of an antibody targeting this modification specifically. Similarly, only a handful of studies so far have investigated the N-terminal (Chen et al., 2009; Mahul-Mellier et al., 2014) and C-terminal (Sano et al., 2021) tyrosine phosphorylations. Here we describe the widespread co-presence of these modifications across LB disorders scattered to neurons and glia. One limitation of our study is that it did not investigate the distribution of ubiquitinated, acetylated or N- and C-terminally truncated aSyn species other than aSyn-120. This is due to the lack of antibodies against this particular modification of the protein, and future efforts to develop appropriate tools to fill this gap will provide an even more complete picture of aSyn heterogeneity. Although the scope of this work primarily covered sporadic and familial LBDs, studies are underway to characterise aSyn pathology in MSA using these tools.

The neuronal seeding model has provided valuable insights into aSyn aggregation and LB-like inclusion formation, and aSyn phosphorylation at Serine 129 has been well described by several groups (Mahul-Mellier et al., 2020, 2018; Volpicelli-Daley et al., 2014, 2011). Until now, the occurrence of other disease-associated PTMs in these models has not been systematically assessed, except for certain C-terminally cleaved aSyn species (Elfarrash et al., 2019; Luk et al., 2012, 2009; Mahul-Mellier et al., 2020, 2018; Volpicelli-Daley et al., 2011). As a result, our understanding of how PTMs influence aSyn seeding and the inclusion formation in these models remains incomplete. In this study, we provide the first comprehensive profiling of post-translationally modified aSyn species in this model. We found that neuronal aSyn accumulations are hyperphosphorylated, and identified the presence of punctate intracellular structures with N- or C-terminal tyrosine phosphorylations (pY39, pY133, pY136) that are not detected by aSyn pS129 antibodies. Exploring the cellular mechanisms underlying these modifications and their impact on aSyn aggregation, aggregate maturation and aSyn-mediated toxicity holds great translational value.

We offer a carefully curated toolset of antibodies that enables a comprehensive assessment of the diverse pathology associated with aSyn. Clearly, not every study design will justify the use of the complete set used here. However, the provided characterisation of the individual antibodies, including their capacities to uncover specific features of aSyn pathology, will enable the choice of appropriate antibodies for specific studies based on solid rationales. In addition, re-visiting the staging studies of PD and other LBDs based on aSyn pathology would greatly benefit from using a good selection of several well-suited antibodies described here to possibly resolve inconsistencies observed when correlating pathology with symptomatology (Jellinger, 2009; Milber et al., 2012; Parkkinen et al., 2008, 2007; Steiner et al., 2018).

By capturing a wide range of aSyn species, we have a valuable opportunity to systematically investigate the correlation between aSyn aggregation, neuronal dysfunction and neurodegeneration in PD and other synucleinopathies. For instance, several studies have reported the absence of LBs in a subset of PD patients with leucine rich repeat kinase 2 (*LRRK2)* mutations (Agin-Liebes et al., 2020; Funayama et al., 2005; Gaig et al., 2007; Giasson et al., 2006; Henderson et al., 2019; Kalia et al., 2015; Marti-Masso et al., 2009; Takanashi et al., 2018; Vilas et al., 2019; Wszolek et al., 2004). By re-examining aSyn pathology using our antibody toolbox, we can conclusively determine whether LBs or other forms of aSyn pathology are genuinely absent in these cases. Additionally, recent findings from our group indicate that certain PTMs, such as O-linked-N-acetylglucosaminylation (O-GlcNAcylation) and nitration, reduce the seeding activity of aSyn fibrils in primary cultures and *in vivo* models (Balana et al., 2023; Donzelli et al., 2023). This suggests that the patterns of PTMs, rather than just the presence of aSyn aggregates, may play a crucial role in determining some of the pathogenic properties of aSyn. Consequently, there is a pressing need for validated tools and methods to accurately map and quantitatively assess the distribution of aSyn PTMs in different brain regions at various stages of disease progression.

In conclusion, we believe that future research aimed at re-visiting 1) PD and other LBD staging in the central nervous system using large cohorts, and 2) aSyn aggregation in the periphery using this validated toolset would open new avenues in our comprehension of spreading of aSyn pathology, and help identify novel therapeutic targets to interfere with these mechanisms.

## MATERIALS AND METHODS

### Generation of aSyn monoclonal antibodies

A combination of aSyn human recombinant proteins and peptides against different forms and modifications of aSyn was used for the immunisation of BALB/c mice (Supplementary Table 1). These were solubilised in PBS and, where appropriate, conjugated to the carrier protein keyhole limpet hemocyanin (KLH). Following pre-immune serum collection, BALB/c mice were subcutaneously injected with the immunogen-complete Freund’s adjuvant (CFA) mixture on Day 0, and with immunogen-incomplete Freund’s adjuvant (IFA) on Day 21 and Day 42. Test bleeds were run on Days 35 and 56, and antibody response was evaluated by ELISA, WB and DB. Animals with strong immunoreactivities were euthanised and their splenocytes surgically harvested.

Hybridoma technology was used for the production of monoclonal antibodies, where the antibody-producing lymphocytes were fused with Sp2/0-Ag14 (ATCC #CRL-8287) mouse myeloma cells using polyethylene glycol (PEG) at a 5:1 ratio. The hybridomas were grown in hypoxanthine-aminopterin-thymidine (HAT) selective media to eliminate unfused myeloma cells. Supernatants of 6 to 44 clones per programme were tested by ELISA, DB and WB against a selected library of aSyn proteins (Supplementary Table 2) to determine the clones with the strongest and the most specific results. Selected clones were further sub-cultured for several rounds to maintain stability, subjected to serial dilution to ensure monoclonality, and screened by ELISA, DB and WB for the identification of positive and specific clones. Isotyping and *in vitro* production of antibodies were carried out with the selected subclones, on an Akta 25 FPLC system using a 25mL protein G sepharose column (Cytiva) according to the instructions of the manufacturer. Briefly, the resin was equilibrated using 10 column volumes (CV) of buffer A (20mM phosphate buffer pH7.2) before loading the filtrated sample on the column. Following a wash step of 10 CV with buffer A, antibodies were eluted using an isocratic elution with buffer B (100mM glycine pH2.7), and were immediately pH-neutralised with 1M Tris buffer pH8.0 upon their harvest in fractionation tubes. Buffer exchange was performed against 10mM PBS using a 30kDa dialysis membrane, and the harvested antibodies were stored at -70 °C. Antibody concentrations were determined using absorbance reading at 280nm, and purity was determined by size exclusion chromatography. An average of 45mg from each of the 12 antibodies were obtained.

### Expression and purification of recombinant proteins

The expression and purification of human and mouse aSyn were carried out as described (Fauvet et al., 2012). Briefly, pT7-7 plasmids encoding variants of mouse and human aSyn were used to transform BL21(DE3) chemically competent *E. coli* and let to grow on an agar dish with ampicillin. One colony was transferred to 400mL of Luria broth (LB) media with ampicillin at 100µg/mL and left to grow at 37 °C on shaker (at 180RPM) for 16h. The small culture was then used to inoculate a large culture of 6L LB media with ampicillin at 100µg/mL. At an optic density (OD_600_) of 0.5-0.6, isopropyl β-D-1-thiogalactopyranoside was added at a final concentration of 1mM to induce aSyn expression. The large culture was left to grow further for 4h on shaker, centrifuged at 4,000*g* for 15min at 4 °C. The pellet was re-suspended on ice in lysis buffer (10mL p/L of culture) containing 20mM Tris pH8.0, 0.3mM phenylmethylsulfonyl fluoride (PMSF) protease inhibitor and cOmplete, mini, EDTA-free protease inhibitor cocktail tablet (Roche #4693159001; one tablet per 10mL lysis buffer). Cells were lysed by sonication (59s-pulse and 59s-no pulse over 5min at 60% amplitude). The lysate was centrifuged at 4 °C for 30min at 20,000*g*, the supernatant boiled for 5-15min at 100 °C, and the centrifugation step repeated. Supernatant was filtered using a 0.22µm syringe filter, and purified via anion exchange chromatography and reverse-phase high performance liquid chromatography (HPLC). The quality control of the proteins was run via analysis by liquid chromatography-mass spectrometry (LC-MS), ultra-performance liquid chromatography (UPLC) and SDS-PAGE separation and Coomassie staining. aSyn nY39, pY39, pS87, pY125 and pS129 protein standards were prepared using semi-synthesis as previously described (Hejjaoui et al., 2012). Recombinant gSyn was purchased from Abcam (#ab48712). Tau 1N4R (Ait-Bouziad et al., 2020), a-beta 42 (Sato et al., 2006) and TDP-43 (Kumar et al., 2023) were expressed and purified as described.

### Generation of aSyn oligomers

Generation of aSyn oligomers was carried out as previously described (Kumar et al., 2020b). Briefly, aSyn human WT monomers were dissolved in PBS for a final concentration of 12mg/mL, and was supplemented with benzonase at 1uL/mL. The solution was incubated in low-protein binding tubes at 37 °C for 5h at 900rpm, centrifuged for 10min at 12,000*g* at 4 °C, and 5mL of supernatant was run through a PBS-equilibrated Hiload 26/600 Superdex 200pg column. Eluted fractions were screened via SDS-PAGE. Oligomeric fractions were characterised by EM, circular dichroism (CD) before being collected, snap frozen and stored at -20 °C.

### Generation of aSyn pre-formed fibrils

Lyophilised human or mouse aSyn WT monomers were re-suspended in PBS for a final concentration of 2-4mg/mL, and the pH was adjusted to 7.5. The protein solution was passed through filters with 100kDa cut-off to remove any spontaneously formed aggregates. Protein concentration was measured via ultraviolet (UV) absorption at 280nm and/or by bicinchoninic acid (BCA) assay. Monomers in solution were left on an orbital shaker (at 1000RPM) for 5 days at 37 °C. For application to cellular seeding models, the fibrils were sonicated to achieve a median fibril length of 50-100nm. The final fibril preparation was characterised for the monomer-to-fibril ratio by sedimentation and filtration assays as described in (Kumar et al., 2020a), for amyloid formation by ThT assay, and for fibril length quantification by electron microscopy analysis.

### Dot/slot blot and Western blot analyses using aSyn recombinant proteins

For the DB analysis, aSyn proteins were diluted in PBS and blotted on a nitrocellulose membrane of 0.22µm in 100µL volume corresponding to 200ng of protein loading (unless indicated otherwise). For the WB analysis, aSyn proteins were diluted in PBS and Laemmli buffer 4X (50% glycerol, 1M Tris at pH 6.8, 20% β-mercaptoethanol, 10% SDS and 0.1% bromophenol blue), loaded onto a 4-16% Tricine gel in 10µL volume corresponding to 100ng of protein loading and transferred onto a nitrocellulose membrane of pore size 0.22µm using a semi-dry transfer system (BioRad) for 45min at 0.5A and 25V. Where appropriate, Ponceau S staining (2% Ponceau S in 5% acetic acid) was applied as protein loading control. The membranes were blocked overnight at 4 °C in Odyssey blocking buffer (Li-Cor). They were incubated with primary antibodies diluted in PBS for 2h at RT, washed three times for 10min in PBS with 0.01% Tween-20 (PBS-T), incubated in dark with secondary antibodies diluted in PBS and washed three times for 10min in PBS-T. For the primary and secondary antibody details, see Supplementary Table 4. The membranes were imaged at 700nm and/or 800nm using the Li-Cor Odyssey CLx imaging system, and the images were processed using Image Studio 5.2.

### Surface plasmon resonance (Biacore)

SPR data were collected on a Biacore 8 K device (GE Healthcare). Antibody (6A3) was immobilised on a CM5 biosensor chip (GE Healthcare) at 10–20 μg/mL concentration in 10 mM acetate solution (GE Healthcare) at pH 4.5 to reach a final surface ligand density of around 2000–4000 response units (RUs). In short, the whole immobilization procedure using solutions of 1-ethyl-3-(3-dimethyl aminopropyl) carbodiimide (EDC) and N-hydroxy succinimide (NHS) mixture, antibody sample and ethanolamine, was carried out at a flow rate of 10 μL/min into the flow cells of the Biacore chip. Firstly, the carboxyl groups on the sensorchip surface were activated by injecting 200 μL of 1:1 (v/v) mixture of EDC/NHS (included in the amine coupling kit, Cytiva Life Sciences) into both flow cells 1 and 2 and followed by the injection of antibodies overflow cell 2 for 180 s. The remaining activated groups in both the flow cells were blocked by injecting 129 μL of 1 M ethanolamine-HCl pH 8.5. The sensor chip coated with antibodies were equilibrated with PBS buffer before the initiation of the binding assays. Serial dilutions of analytes such as α-syn monomers (human WT 1-20 or human WT 1-140) at a concentration ranging between 2.5 μM to 0.1 μM in PBS buffer were injected into both flow cells at a flow rate of 30 μL/min at 25 °C. Each sample cycle has the contact time (association phase) of 120 s and is followed by a dissociation time of 600 s. After every injection cycle, surface regeneration of the Biacore chips was performed using 10 mM glycine (pH 3.0).

### Mouse primary neuronal culture and seeding assay

Primary hippocampal and cortical neurons were collected from P0 pups of WT (C57BL/6J-RccHsd, Harlan) or aSyn KO (C57BL/6J-OlaHsd, Harlan) mice, according to the dissection procedure described elsewhere (Steiner et al., 2002). Following the plating in poly-L-lysine-coated plates (300,000 cells/mL), the neurons were left to mature at 37 °C with 5% CO_2_. WT neurons were treated with 70nM aSyn mouse WT PFFs on day *in vitro* (DIV)7 and left to incubate for 14 days; and aSyn KO neurons on DIV20 and left to incubate for 24h, as described (Mahul-Mellier et al., 2020, 2018; Volpicelli-Daley et al., 2014, 2011).

### Immunocytochemistry and confocal imaging

Mouse primary neurons were washed twice in PBS, fixed with 4% PFA for 20min at RT and stained as described elsewhere (Mahul-Mellier et al., 2015). The antibodies used for ICC are detailed in Supplementary Table 4. Imaging was carried out on a confocal laser-scanning microscope (LSM 700, Carl Zeiss, Germany) and image analysis on Zen Digital Imaging software (RRID: SCR_013672).

### Cell lysis, sequential extraction and Western blotting

Mouse primary neurons were washed in PBS twice on ice and extracted in Tris-buffered saline (TBS; 50mM Tris and 150mM NaCl at pH7.5) with 1% Triton X-100 (Tx-100) and with protease inhibitor (PI) cocktail (1:100), 1mM phenylmethane sulfonyl fluoride (PMSF), phosphatase inhibitor cocktails 2 and 3 (1:100), as described previously (Mahul-Mellier et al., 2020, 2018; Volpicelli-Daley et al., 2014, 2011). The cells were lysed via sonication (1s intermittent pulse 10 times at 20% amplitude) using a small probe (Sonic Vibra Cell, Blanc Labo, Switzerland). The lysate was incubated for 30min on ice and centrifuged at 100,000*g* for 30min at 4 °C. The supernatant i.e. the ‘soluble’ fraction, was collected and diluted in 4X Laemmli buffer (50% glycerol, 1M Tris at pH 6.8, 20% β-mercaptoethanol, 10% SDS and 0.1% bromophenol blue). As the wash step, the pellet was re-suspended in lysis buffer, sonicated and centrifuged as described above. Supernatant was discarded and the pellet re-suspended in TBS with 2% sodium dodecyl sulphate supplemented with protease and phosphatase inhibitors as described above. The re-suspension, i.e. the ‘insoluble fraction’, was sonicated (1s intermittent pulse 15 times at 20% amplitude) and diluted in 4X Laemmli buffer. Protein concentration was determined by BCA assay separately for the soluble and the insoluble fractions.

The soluble and insoluble fractions were separated on a 16% Tricine gel (ThermoFisher), transferred onto a nitrocellulose membrane of pore size 0.22µm using a semi-dry transfer system (BioRad) for 45min at 0.5A and 25V. The membranes were blocked overnight at 4 °C in Odyssey blocking buffer (Li-Cor) and washed three times for 10min in PBS with 0.01% Tween-20 (PBS-T). Membranes were incubated with primary antibodies diluted in PBS for 2h at RT, washed three times for 10min in PBS-T, incubated in dark with secondary antibodies diluted in PBS and washed three times for 10min in PBS-T. For the antibody details, see Supplementary Table 4. The membranes were imaged at 700nm and/or 800nm using the Li-Cor Odyssey Calyx imaging system, and the images processed using Image Studio 5.2.

### Animals and intra-striatal stereotaxic injection procedure

All animal experimentation was performed in compliance with the European Communities Council Directive of 24 November 1986 (86/609EEC) and with approval of the Cantonal Veterinary Authorities (Vaud, Switzerland) and the Swiss Federal Veterinary Office (authorisation number VD2067.2). C57BL/6JRj male mice (Janvier Labs) at 3 months of age were stereotaxically injected with aSyn mouse WT PFFs (5mg in 2mL PBS) in the right dorsal striatum. 6 months post-injection, the animals were sacrificed by intracardiac perfusion with heparinised sodium chloride (NaCl; 0.9%), and fixed with 4% paraformaldehyde (PFA) in PBS overnight, and paraffin-embedded for immunohistochemical studies. For the antibody validation studies, naïve adult aSyn KO mice (C57BL/6J OlaHsd, Harlan) were sacrificed and brain sections were prepared in the same way as for PFF-injected WT animals.

### Immunofluorescent labelling and imaging of mouse brain tissue

WT and aSyn KO mouse paraffin-embedded brain sections were cut coronally to 4µm and dewaxed. Epitope retrieval was carried out in 10mM trisodium citrate buffer at pH6.0 for 20min at 95 °C. Tissue blocking was run by incubation in 3% bovine serum albumin (BSA) and 0.1% Tx-100 in PBS for 1h at RT. Sections were then incubated with the primary antibody solution overnight at 4 °C, and with the secondary antibody solution for 60min at RT (Supplementary Table 4). Slides were mounted using an aqueous mounting medium, and tiled imaging was carried out on the Olympus VS120 microscope.

### Human brain tissue samples

The cases selected from the Queen Square Brain Bank (QSBB), UCL Institute of Neurology in London, UK, were collected in accordance with the London Multicentre Research Ethics Committee-approved protocol. All donors had given written informed consent prior to the brain collection. The samples were stored under the licence (no: 12198) approved by the Human Tissue Authority. The ethical approval for research was given by the National Research Ethics Service (NRES) Committee London Central. Case demographics are detailed in Table 5.

**Table 5:** Demographics of the human post-mortem cases included in this study. AD = Alzheimer’s disease; CBD = corticobasal degeneration; CTR = control; DLB = dementia with Lewy bodies; f = female; FTLD-TDP/C = frontotemporal lobar degeneration of transactive response DNA-binding protein 43 type C; h = hours; ID = identity; LB = Lewy body; m = male; na = not available; PD = Parkinson’s disease; PDD = Parkinson’s disease with dementia; PiD = Pick’s disease; PSP = progressive supranuclear palsy; y = years

| case ID | gender (m/f) | age at diagnosis (y) | age at death (y) | disease duration (y) | postmortem delay (h) | Braak stage (LB-type pathology) | Braak stage (AD-type pathology) |
| --- | --- | --- | --- | --- | --- | --- | --- |
| PD1 | f | 65 | 86 | 21.0 | 78.4 | 6 | 1 |
| PD2 | m | 55 | 75 | 20.0 | 88.5 | 6 | 1 |
| PD3 | m | 60 | 84 | 24.0 | 71.0 | 6 | 2 |
| PD4 | m | 62 | 80 | 18.5 | 67.0 | 6 | 3 |
| PD5 | m | 68 | 80 | 11.9 | 100.0 | 6 | 2 |
| PD6 | f | 61 | 81 | 20.0 | 135.0 | 6 | 2 |
| PD7 | m | 80 | 89 | 8.5 | 26.0 | 6 | 4 |
| PDD1 | m | 77 | 92 | 14.0 | 41.0 | 6 | 3 |
| PDD2 | m | 61 | 80 | 19.0 | 66.0 | 6 | 2 |
| DLB1 | f | 78 | 89 | 11.4 | 63.0 | 6 | 3 |
| SNCA G51D1 | m | 19 | 49 | 30.0 | 43.0 | na | na |
| SNCA G51D2 | f | 69 | 75 | 6.5 | 62.0 | na | 1 |
| SNCA G51D3 | m | 45 | 52 | 6.5 | 85.3 | na | 1 |
| SNCA H50Q1 | f | 70 | 83 | 12.5 | na | na | na |
| SNCA duplication1 | m | 55 | 62 | 6.9 | 143.0 | 6 | 2 |
| AD1 | m | 54 | 64 | 9.8 | 95.5 | 0 | 6 |
| AD2 | f | 48 | 64 | 16.6 | 70.0 | na | 6 |
| PiD1 | m | 55 | 71 | 16.4 | 51.0 | 0 | 0 |
| PiD2 | m | 52 | 65 | 13.5 | 48.0 | 0 | 0 |
| FTLD-TDP/C1 | f | 51 | 65 | 14.0 | 22.0 | 0 | 0 |
| FTLD-TDP/C2 | m | 53 | 71 | 17.3 | 54.0 | 0 | 0 |
| PSP1 | f | 74 | 79 | 6.3 | 105.0 | 0 | 2 |
| PSP2 | m | 68 | 75 | 7.0 | 57.0 | 0 | 2 |
| CBD1 | m | 51 | 60 | 9.0 | 40.0 | na | 0 |
| CBD2 | m | 60 | 65 | 5.0 | 50.0 | 0 | 2 |
| CTR1 | f | 67 | 84 | 16.0 | 40.0 | 0 | 2 |
| CTR2 | m | 63 | 80 | 16.6 | 11.0 | 0 | 2 |
| CTR3 | f | na | 89 | na | 24 | 0 | 2 |
| CTR4 | m | na | 80 | na | 48 | 0 | 2 |

### Immunohistochemistry of human brain samples with 3,3′-diaminobenzidine (DAB) revelation and imaging

FFPE sections were cut sequentially to 8µm of thickness and de-paraffinised. The rationale for the selection of the final epitope retrieval approach and dilution for each antibody was to enable the antibodies to reveal the optimal number of pathological inclusions with minimal non-specific background. For epitope retrieval, the sections were treated with 80-100% formic acid for 10min at room temperature (RT) and/or with citrate buffer (pH6.0) for 10min at 121°C under pressure. Sections were treated with 3% hydrogen peroxide in PBS for 30min to quench the endogenous peroxidase. After the blocking in 10% foetal bovine serum (FBS) for 30min, sections were incubated in primary antibody solution overnight at 4°C. For the primary antibody details and their optimised IHC settings, see Supplementary Tables 4-5. After being rinsed in PBS-Tween 0.1% (PBS-T), sections were incubated in the secondary antibody-horseradish peroxidase (HRP) complex from the REAL EnVision detection kit (Dako #K5007) for 1h at RT. Sections were rinsed in PBS-T before visualisation with 3,3’-diaminobenzidine (DAB). They were counterstained with haematoxylin, cleared in xylene and mounted using distyrene plasticiser xylene (DPX).

### Immunofluorescent labelling and imaging of human brain tissue

Following the blocking in 3% BSA and 0.3% Tx-100 in PBS for 60 min at RT, sections were washed in PBS for 5 min and incubated for 1 min in TrueBlack lipofuscin autofluorescence quencher (Biotium #23,007) in 70% ethanol. The sections were washed in PBS (3×5 min) and incubated in primary antibodies overnight at 4 °C. For the primary antibody details and their optimised IF settings, see Supplementary Tables 4-5. After rinsing in PBS, the sections were incubated in secondary antibodies for 1 h at RT in dark and washed in PBS. Where applicable, the sections were then incubated in biotinylated primary antibody solution at 4 °C overnight, rinsed in PBS, incubated in streptavidin secondary antibody solution for 1h at RT in dark and rinsed in PBS. The slides were mounted using an aqueous mounting medium with DAPI (Vector Laboratories #H-1500-10). Imaging was carried out on DM5500 B upright microscope (Leica, Germany), and image analysis on Leica Application Suite X (RRID:SCR_013673).

## FIGURE LEGENDS

**Supplementary Figure 1:** Specificity validation of the novel aSyn PTM antibodies using human recombinant aSyn standards. Novel monoclonal aSyn PTM antibodies were validated by **(A)** WB screening. The 5E1-G8 antibody showed non-specific positivity to aSyn WT by WB (red arrow). Further **(B)** DB and **(C)** WB analyses on 6A3-E9 showed that this antibody is specific to human aSyn truncated at residue 120. **(D)** SPR sensograms showed the binding responses of immobilised antibody 6A3-E9 against varying concentrations of aSyn human 1-120 (top) or aSyn human WT (bottom). aSyn = alpha-synuclein; CTR = control; DB = dot/slot blot; PTM = post-translational modification; SPR = surface plasmon resonance; WB = Western blot; WT = wild-type

**Supplementary Figure 2:** aSyn antibody sensitivity to neighbouring aSyn PTMs, reactivity to other synuclein family proteins and amyloidogenic proteins. **(A)** The amino acid sequence of aSyn human and its mouse orthologue. **(B)** The synuclein family comprises three homologous proteins – aSyn, bSyn and gSyn. The residue differences are highlighted in red. **(C)** The sensitivities of non-modified aSyn antibodies to PTMs neighbouring or overlapping their epitopes were explored by DB (red arrows). Protein loading control was run via Ponceau S staining. **(D)** Reactivity of the non-modified aSyn antibodies to other members of the synuclein family and to other amyloidogenic proteins tau (1N4R), a-beta 42 and TDP-43 were studied by DB. Protein loading control was run via Ponceau S staining. Blue arrows indicate reactivity to bSyn, green arrows to gSyn and red arrows cross-reactivity to other amyloidogenic proteins. a-beta = amyloid-beta; aSyn = alpha-synuclein; bSyn = beta-synuclein; DB = dot/slot blot; gSyn = gamma-synuclein; h = human; IB = immunoblot; m = mouse; NAC = non-amyloid component; PBS = phosphate buffered saline; PTM = post-translational modification; tau = tubulin-associated unit; TDP-43 = transactive response DNA-binding protein 43kDa; WT = wild-type

**Supplementary Figure 3:** Specificity validation of non-modified aSyn antibodies on aSyn KO hippocampal and cortical neurons by ICC. PBS- and PFF-treated aSyn KO **(A)** hippocampal and **(B)** cortical neurons were immunostained to validate the specificity of the antibodies with epitopes against the N-terminus, NAC region and C-terminus of aSyn. Blue arrows indicate bSyn positivity in PBS-treated neurons, green arrows indicate positivity to aSyn mouse WT fibrils in PFF-treated neurons, and red arrows indicate non-specific background both in PBS- and PFF-treated neurons. aSyn = alpha-synuclein; bSyn = beta-synuclein; DAPI = 4’,6-diamidino-2-phenylindole; ICC= immunocytochemistry; KO = knockout; NAC = non-amyloid component; MAP2 = microtubule-associated protein 2; PBS = phosphate buffered saline; PFF = pre-formed fibril; WT = wild-type

**Supplementary Figure 4:** Specificity validation of aSyn PTM antibodies on aSyn KO hippocampal and cortical neurons by ICC. PBS- and PFF-treated aSyn KO **(A)** hippocampal and **(B)** cortical neurons were immunostained to validate the specificity of the antibodies with epitopes against the PTMs of aSyn. Green arrows indicate positivity to aSyn mouse WT fibrils in PFF-treated neurons, and red arrows indicate non-specific background both in PBS- and PFF-treated neurons. aSyn = alpha-synuclein; bSyn = beta-synuclein; DAPI = 4’,6-diamidino-2-phenylindole; ICC = immunocytochemistry; KO = knockout; MAP2 = microtubule-associated protein 2; PBS = phosphate buffered saline; PFF = pre-formed fibril; PTM = post-translational modification; WT = wild-type

**Supplementary Figure 5:** Specificity validation of aSyn antibodies on aSyn KO hippocampal and cortical neurons by WB. PBS- and PFF-treated aSyn KO hippocampal and cortical neurons were separated to soluble and insoluble fractions by sequential extraction and stained using **(A)** non-modified and **(B)** aSyn PTM antibodies for specificity validation. Green arrows indicate bands specific to aSyn mouse WT fibrils, blue arrows indicate bSyn-specific bands, and red arrows indicate non-specific background. aSyn = alpha-synuclein; bSyn = beta-synuclein; ctx = cortex; hipp = hippocampus; IB = immunoblot; NAC = non-amyloid component; KO = knockout; PBS = phosphate buffered saline; PFF = pre-formed fibril; PTM = post-translational modification; WB = Western blot; WT = wild-type

**Supplementary Figure 6:** Specificity validation of aSyn antibodies on aSyn KO mouse brain tissue by IF. The in-house and commercial antibodies against **(A)** non-modified aSyn and **(B)** aSyn PTMs were screened for their specificity using aSyn KO mouse amygdala sections. Red arrows indicate non-specific background. aSyn = alpha-synuclein; DAPI = 4’,6-diamidino-2-phenylindole; IF = immunofluorescence; KO = knockout; PTM = post-translational modification

**Supplementary Figure 7:** IF labelling of PD cingulate cortex using the aSyn N-terminal LASH-EGTNter 1-20 or LASH-BL 34-45, aSyn C-terminal BL 4B12 103-108 or AB 134-138, and aSyn pS129 BL 81A-biotin antibodies. LBs are marked with asterisks, and LNs with arrows. Representative images from PD1 cingulate cortex are taken using Leica DM5500 B upright microscope at 20x magnification. aSyn = alpha-synuclein; DAPI = 4’,6-diamidino-2-phenylindole; IF = immunofluorescence; IHC = immunohistochemistry; LB = Lewy body; LN = Lewy neurite; PD = Parkinson’s disease

**Supplementary Figure 8:** Specificity validation of aSyn antibodies on human post-mortem tissues via IHC. The frontal cortices of PSP and CBD, and the hippocampi and the EC of AD, PiD and FTLD-TDP/C were stained using the selection of aSyn non-modified and PTM antibodies. Cross-reactivity was not observed with pathological accumulations positive for a-beta (antibody: Agilent anti-a-beta), p-tau (antibody: TF AT8) or TDP-43 (antibody: LSBio 6F/3D). Arrows indicate aSyn-positive structures detected on each tissue. Representative images taken from the entorhinal cortices (layers V-VI) and hippocampi (dentate gyrus) of AD1-2, PiD1-2 and FTLP-TDP/C1-2, and the frontal cortices (layers V– VI) of PSP1-2 and CBD1-2. a-beta = amyloid-beta; AD = Alzheimer’s disease; aSyn = alpha-synuclein; CBD = corticobasal degeneration; ctx = cortex; EC = entorhinal cortex; FTLD-TDP/C = frontotemporal lobar degeneration of transactive response DNA-binding protein 43 type C; hipp = hippocampus; IHC = immunohistochemistry; PiD = Pick’s disease; PSP = posterior supranuclear palsy; p-tau = phosphorylated tubulin-associated unit; PTM = post-translational modification; TDP-43 = transactive response DNA binding protein 43kDa

**Supplementary Figure 9:** Complementary images to Figure 5B-C. Staining of the (A) SN and (B) cingulate cortex (layers V–VI) of healthy controls with aSyn antibodies. aSyn = alpha-synuclein; CTR = control; SN = substantia nigra

**Supplementary Table 1:** A list of all the programmes initialised to generate monoclonal antibodies against aSyn. aSyn = alpha-synuclein; FL = full-length; KLH = keyhole limpet hemocyanin; na = not available

**Supplementary Table 2:** A list of all aSyn proteins and peptides included in this study. aSyn = alpha-synuclein; Da = dalton; FL = full-length; h = human; m = mouse; Mw = molecular weight

**Supplementary Table 3:** A list of all the purified antibodies and their epitopes after finalisation of the monoclonal antibody generation programmes. aSyn = alpha-synuclein; hu = human; IgG = immunoglobulin G; K = kappa; mc = monoclonal; mus = mouse

**Supplementary Table 4:** A list of all the primary and secondary antibodies used in this study. a-beta = amyloid-beta; aSyn = alpha-synuclein; ch = chicken; h = human; MAP2 = microtubule-associated protein 2; mc = monoclonal; mus or m = mouse; pc = polyclonal; rab or r = rabbit; tau = tubulin-associated unit; TDP-43 = transactive response DNA-binding protein 43kDa

**Supplementary Table 5:** Optimised IHC settings for the antibodies used on human post-mortem brain tissues. *AC+FA pre-treatment was applied in IF studies for all antibodies. a-beta = amyloid-beta; AC = autoclave; aSyn = alpha-synuclein; FA = formic acid; IF = immunofluorescence; IHC = immunohistochemistry; na = not applicable; tau = tubulin associated unit; TDP-43 = transactive response DNA-binding protein 43kDa

### Contributions of the authors

HAL conceived and conceptualised the study. HAL and MFA designed the experiments. STK ran the SPR experiment. JB provided the aSyn KO and PFF-seeded WT mouse brain tissues for IHC. SJ generated and characterised the aSyn oligomers. JLH selected the human cases. LP contributed to the optimisation of the IHC assays on human cases. JLH, YM and LP contributed to the analysis of the human data. MFA performed all other experiments involved in the study and wrote the first draft of the manuscript, which was reviewed by all the authors.

HAL: Hilal A. Lashuel

JB: Johannes Burtscher

JLH: Janice L. Holton

LP: Laura Parkkinen

MFA: Melek Firat Altay

SJ: Somanath Jagannath

STK: Senthil T. Kumar

YM: Yasuo Miki

## Acknowledgments

This work was supported by grants from the Swiss National Science Foundation (31ER30_186198), Michael J Fox Foundation (MJFF-020698) and EPFL. The Queen Square Brain Bank is supported by the Reta Lila Weston Institute of Neurological Studies, UCL Queen Square Institute of Neurology. We thank Jonathan Ricci and Driss Boudeffa for contributing to the Western blot and dot blot analysis experiments during the antibody generation stage. We thank Dr Sergey Nazarov for providing us with the aSyn fibril reconstruction images in Figure 1B, and Dr Galina Limorenko and Lixin Yang with tau, a-beta and TDP-43 recombinant proteins. We thank Dr Galina Limorenko also for her valuable feedback towards finalising the manuscript. We thank all the individuals and their families for agreeing to donate to QSBB and OBB.

