## Supplementary data for "Development and validation of an expanded antibody toolset that captures alpha-synuclein pathological diversity in Lewy body diseases"

### SUPPLEMENTARY FIGURE 1

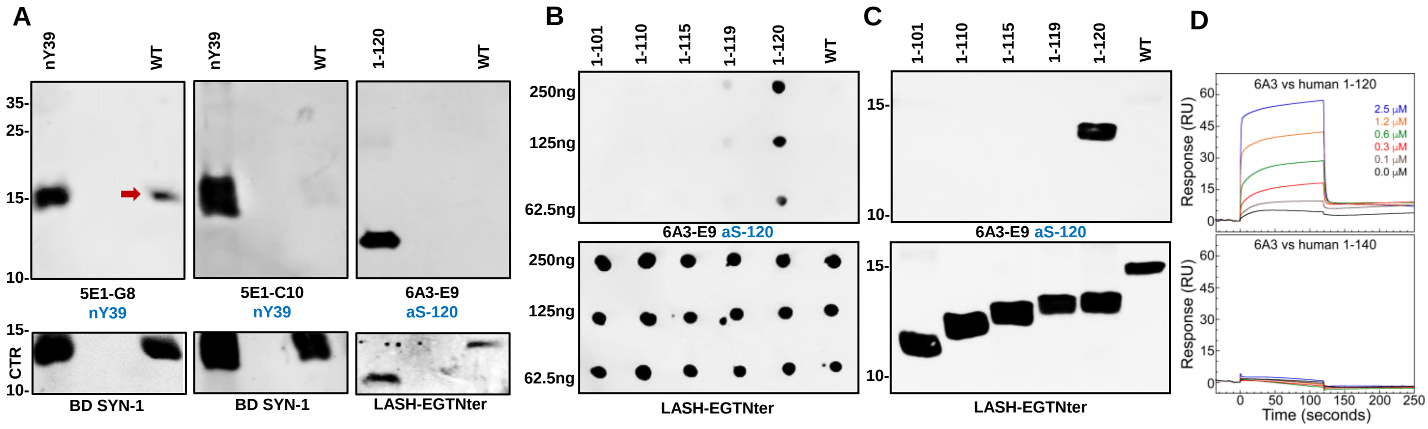

### SUPPLEMENTARY FIGURE 2

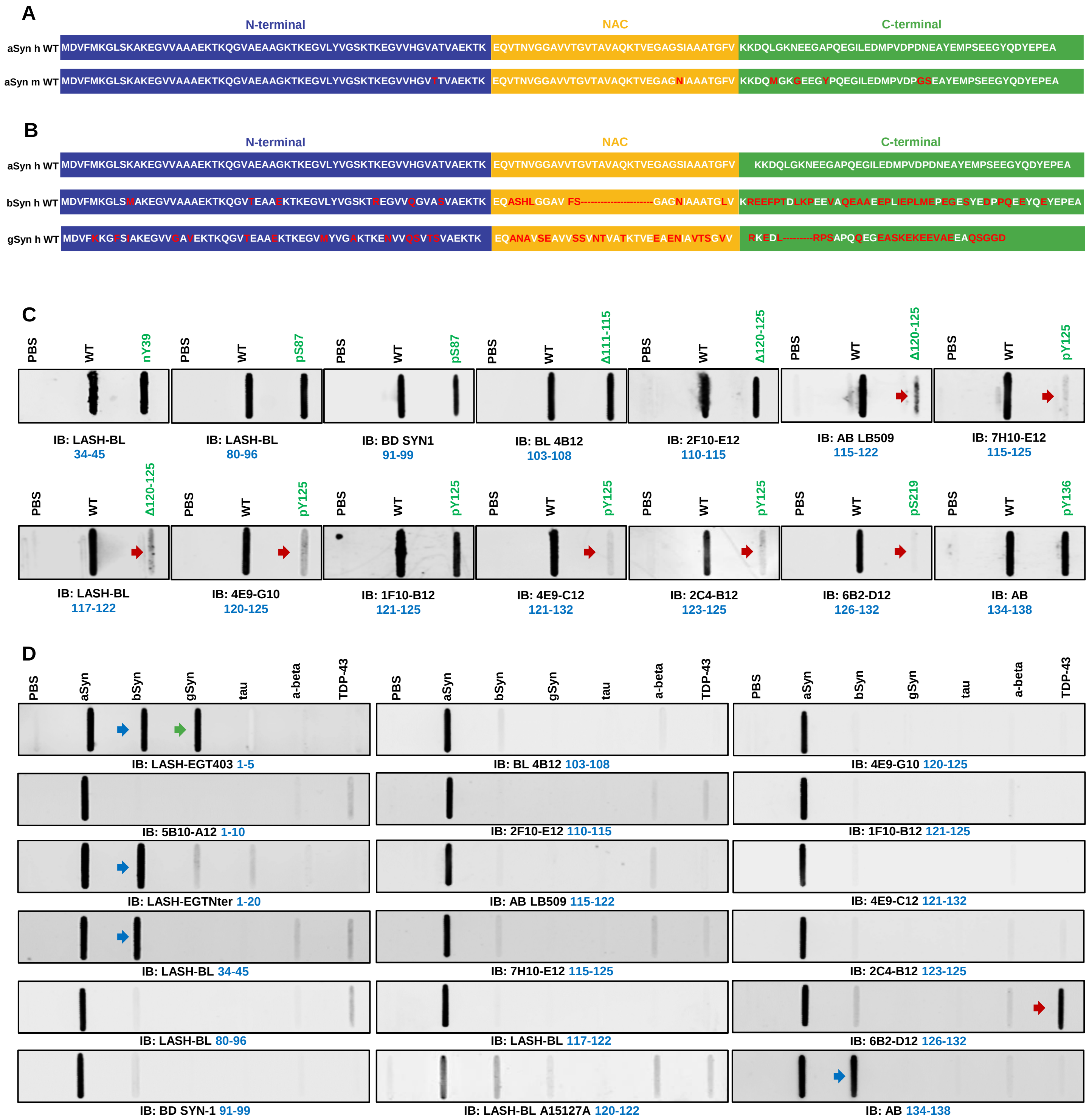

### SUPPLEMENTARY FIGURE 3

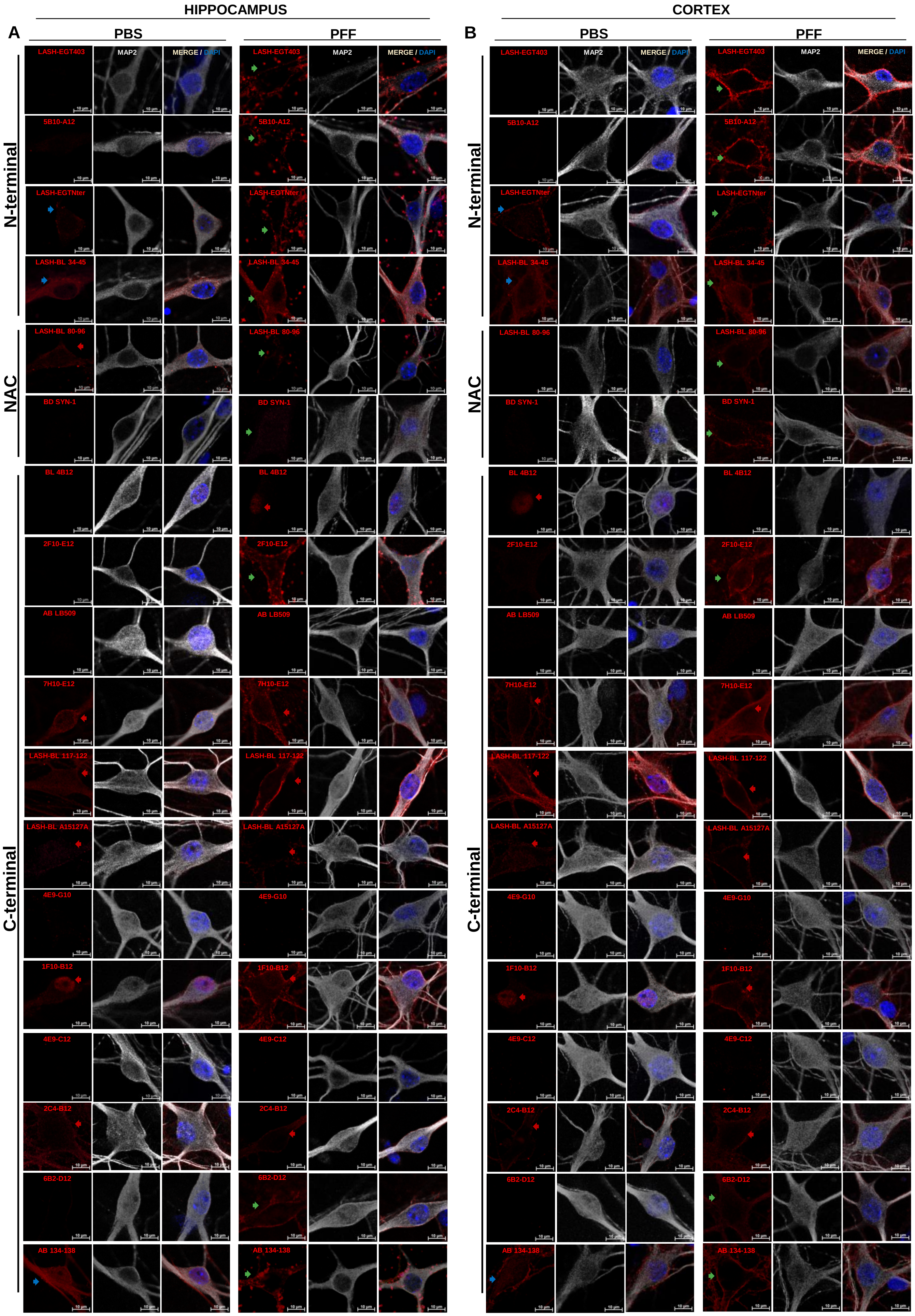

#### SUPPLEMENTARY FIGURE 4

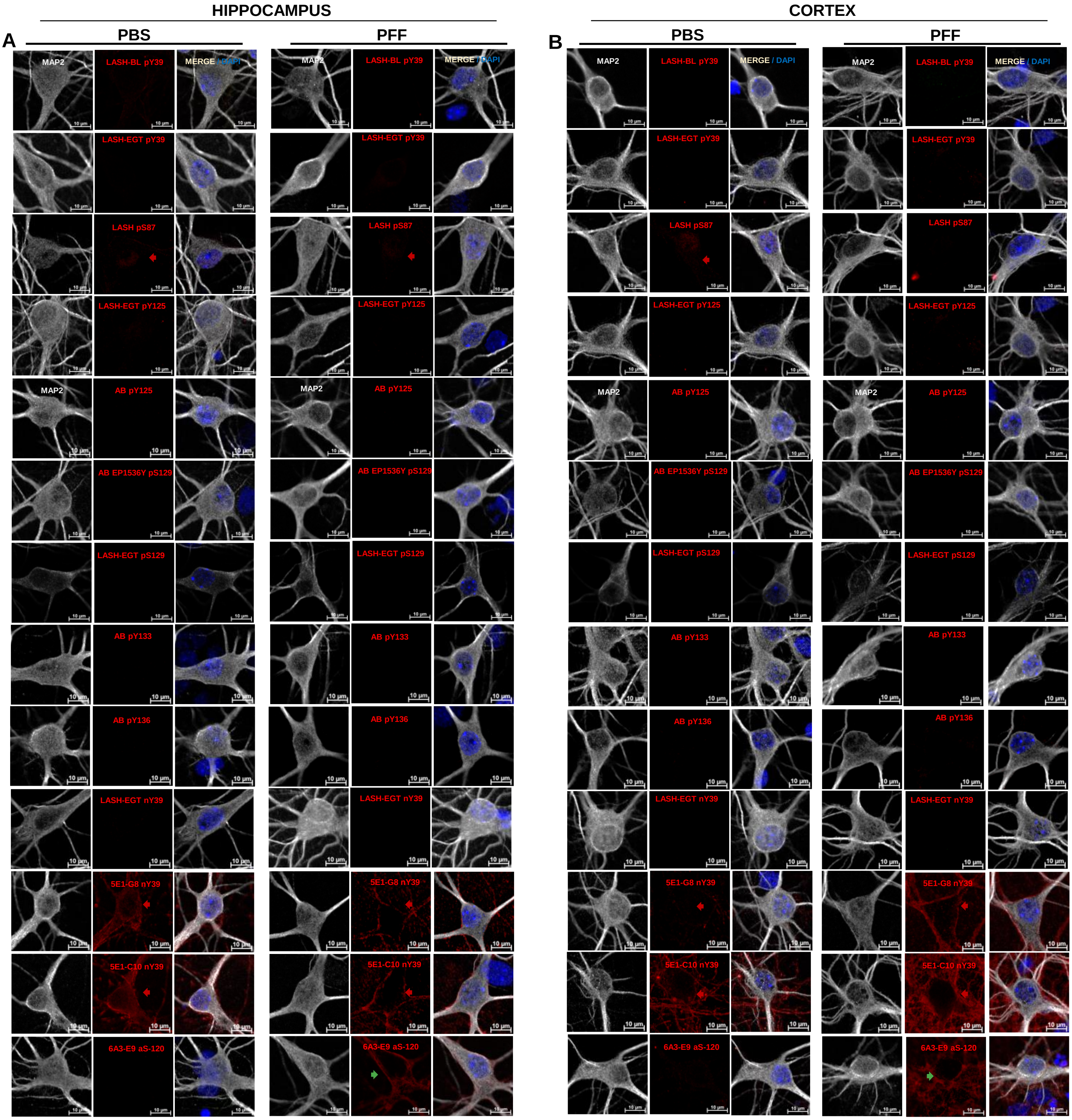

### SUPPLEMENTARY FIGURE 5

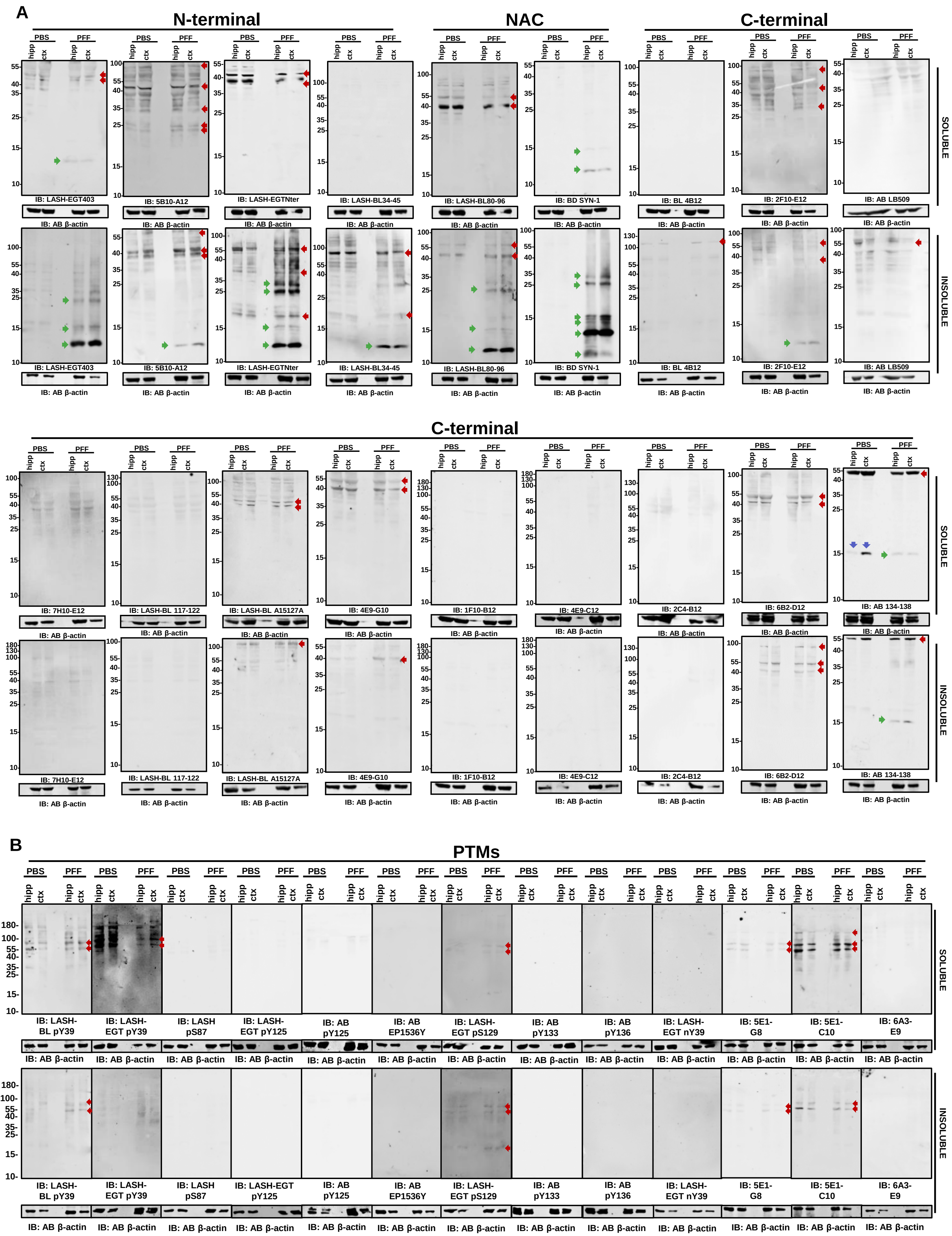

SUPPLEMENTARY FIGURE 6

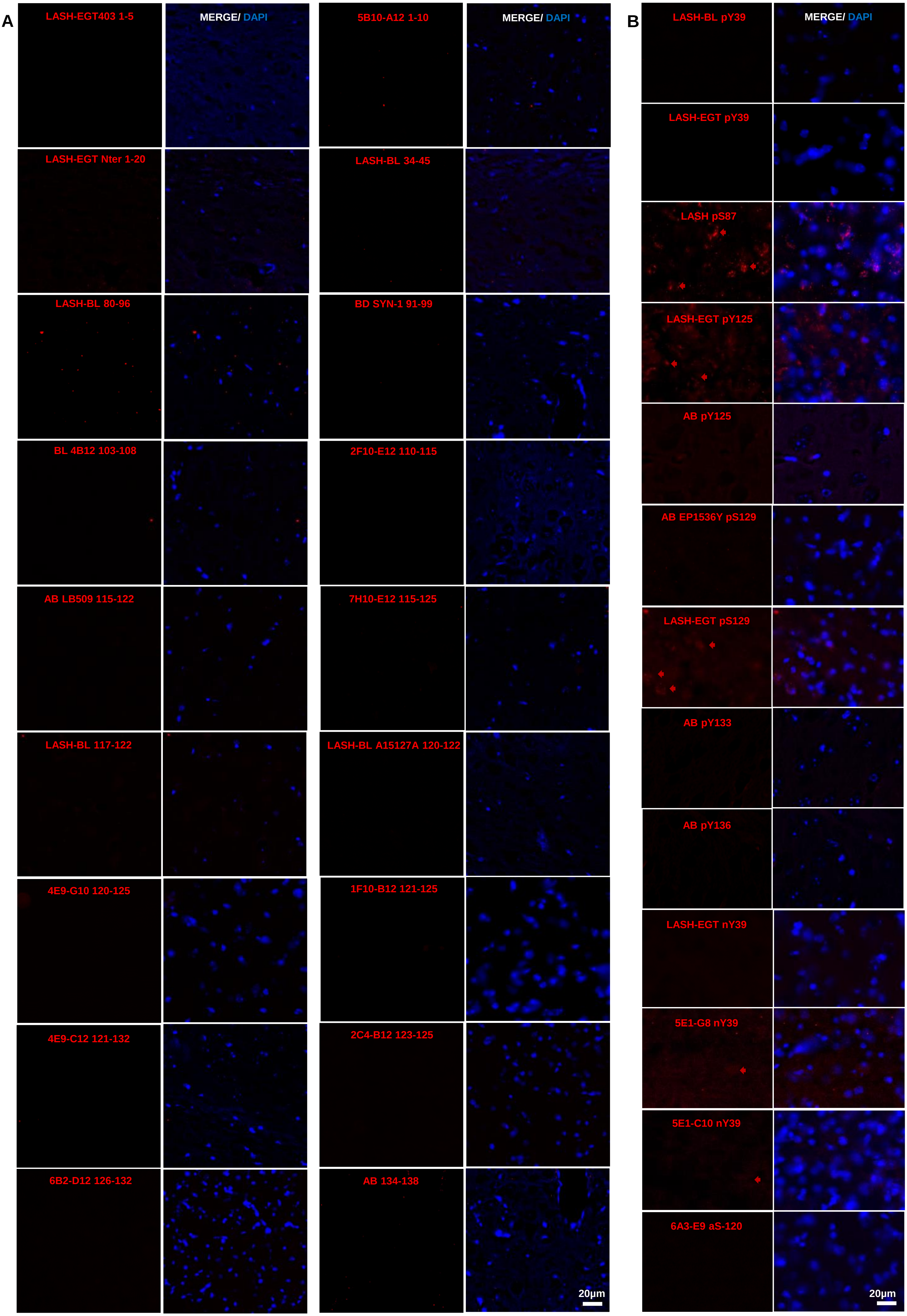

#### SUPPLEMENTARY FIGURE 7

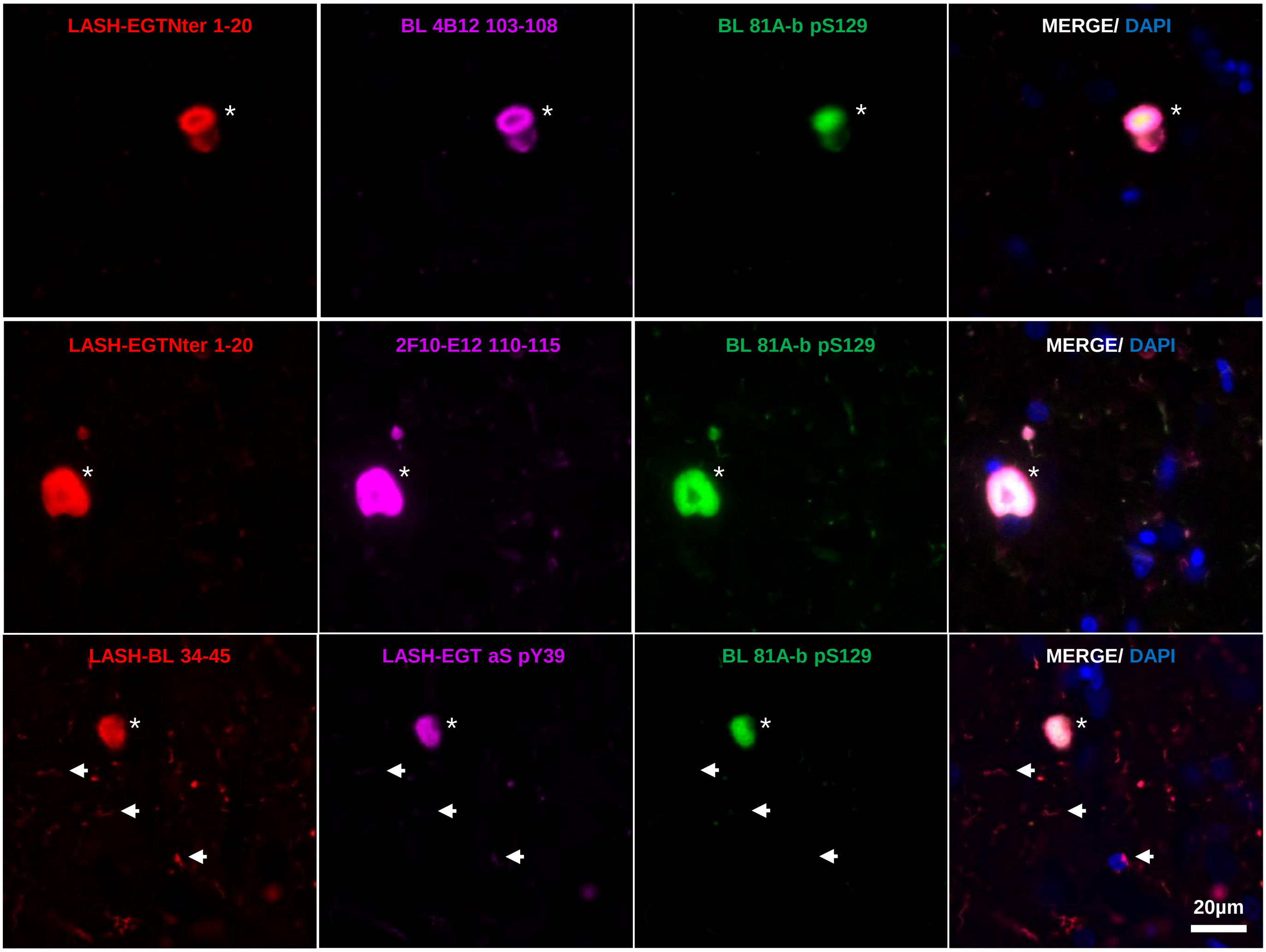

### SUPPLEMENTARY FIGURE 8

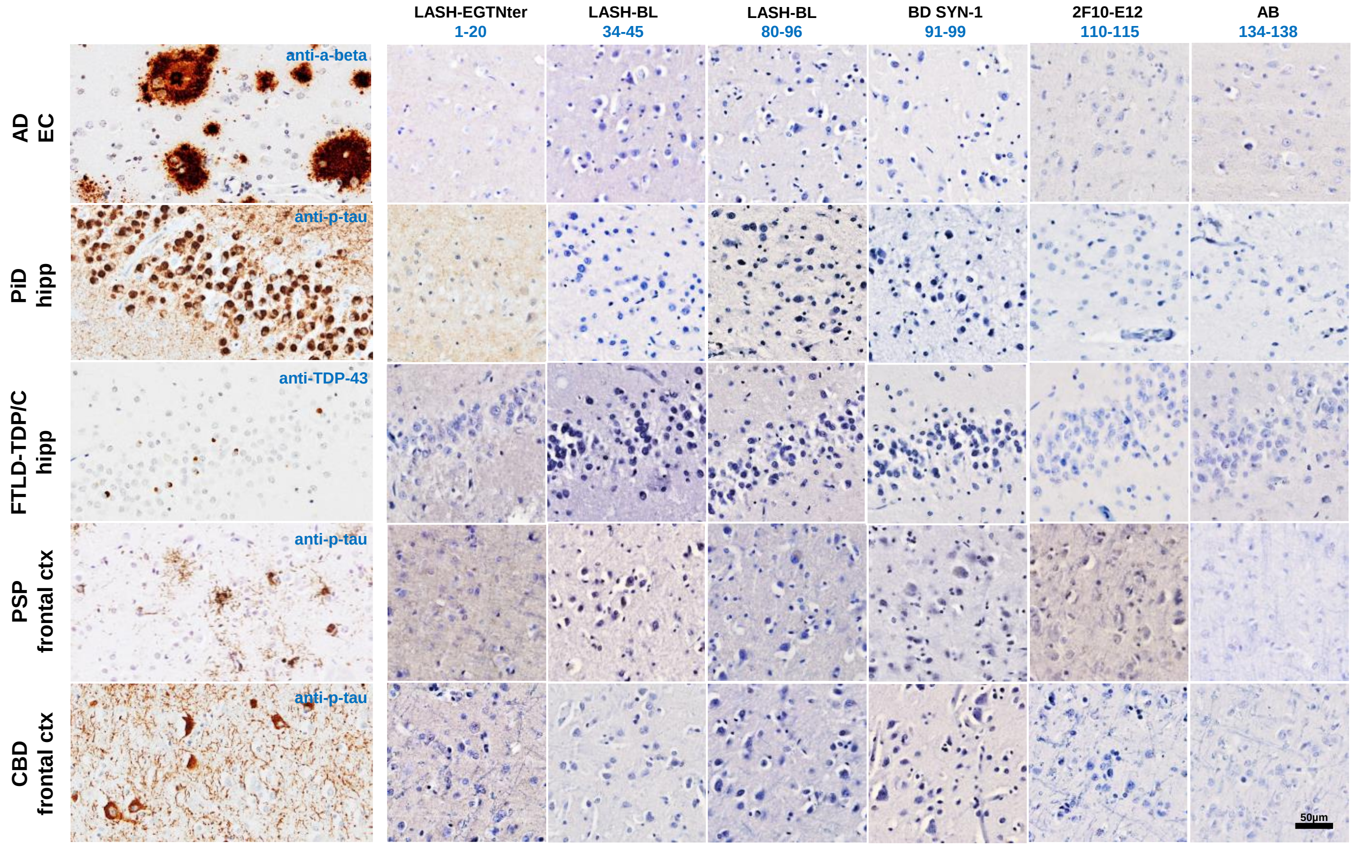


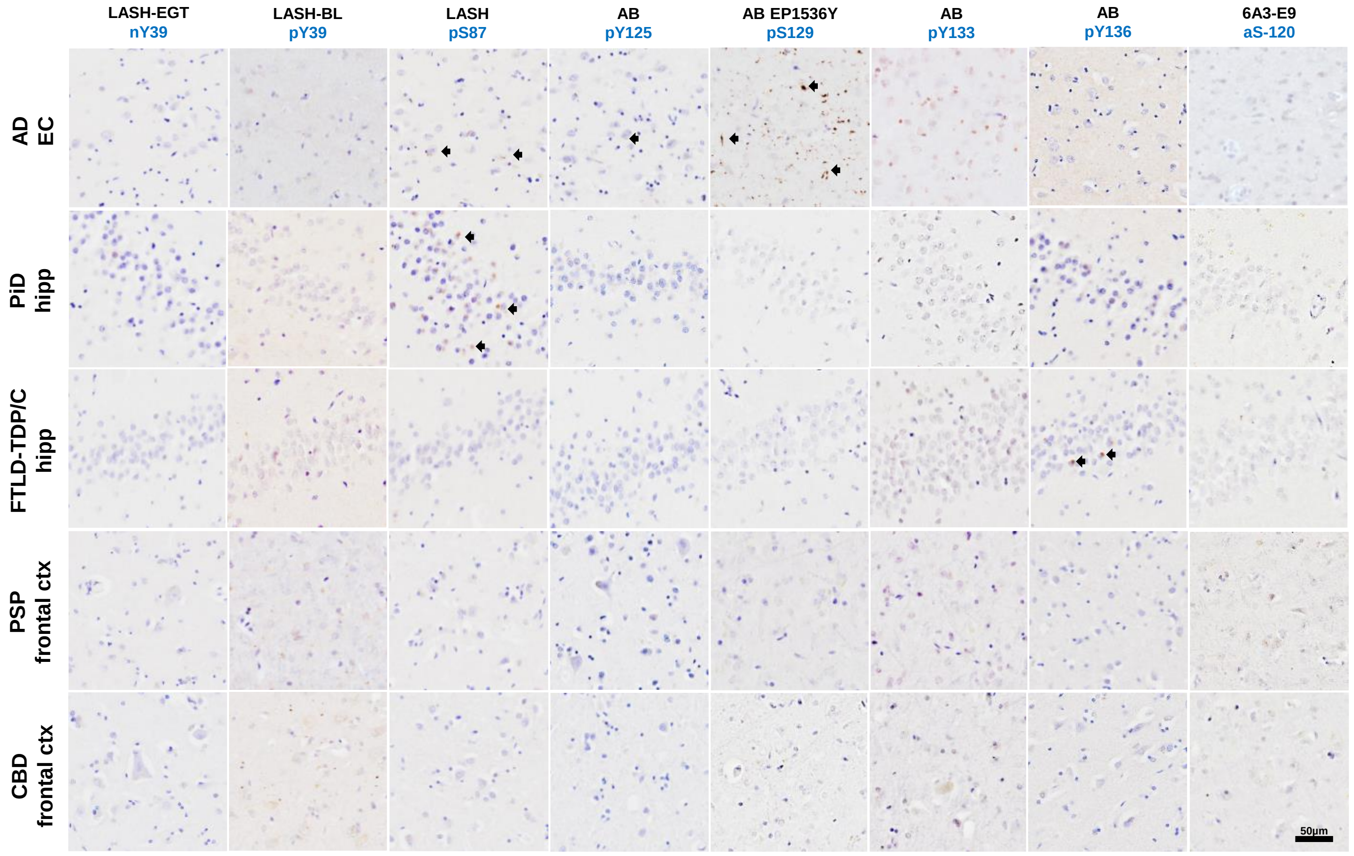

### SUPPLEMENTARY FIGURE 9

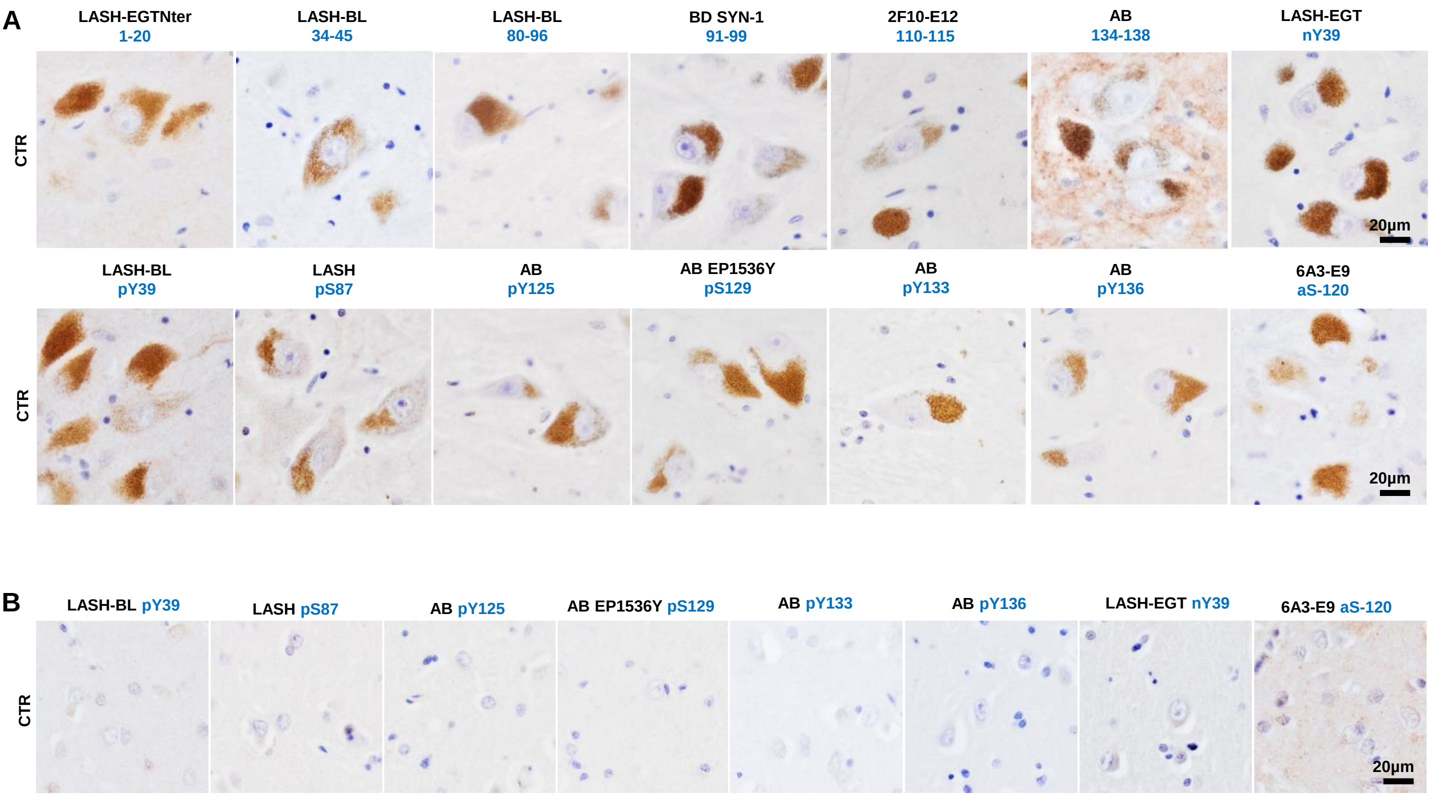

**Supplementary Table 1:** A list of all the programmes initialised to generate monoclonal antibodies against aSyn.

| programme | immunogen | carrier | antigen sequence |
|---|---|---|---|
| LASH-EGT403 | UniProtKB - P37840 (SYUA_HUMAN) **1-20** | KLH | MDVFMKGLSK AKEGVVAAAE |
| LASH-EGT404 | UniProtKB - P37840 (SYUA_HUMAN) **1-140** | na | recombinant aSyn human FL (1-140) |
| LASH-EGT405 | UniProtKB - P37840 (SYUA_HUMAN) **113-127** | KLH | LEDMPVDP DNEAYEM |
| LASH-EGT406 | UniProtKB - P37840 (SYUA_HUMAN) **108-120** | KLH | PQE GILEDMPVDP |
| LASH-EGT407 | UniProtKB - P37840 (SYUA_HUMAN) **124-135 pS129** | KLH | AYEMP(pS)E EGYQD |
| LASH-EGT408 | UniProtKB - P37840 (SYUA_HUMAN) **108-140** | na | PQE GILEDMPVDP DNEAYEMPSE EGYQDYEPEA |
| LASH-EGT409 | UniProtKB - P37840 (SYUA_HUMAN) **120-135 pY125/pS129** | KLH | P DNEA(pY)EM P(pS)E EGYQD |
| LASH-EGT410 | UniProtKB - P37840 (SYUA_HUMAN) **1-119** | na | recombinant aSyn human 1-119 |
| LASH-EGT416 | UniProtKB - P37840 (SYUA_HUMAN) **34-45 nY39** | KLH | KEGVL(nY)VGSKTK |

aSyn = alpha-synuclein; FL = full-length; KLH = keyhole limpet hemocyanin; na = not available

**Supplementary Table 2:** A list of all aSyn proteins and peptides included in this study.

| aSyn protein/ peptide | Mw/ Da | aSyn protein/ peptide | Mw/ Da | aSyn protein/ peptide | Mw/ Da |
|---|---|---|---|---|---|
| h/m 1-11 | 1,227 | h 39-140 | 10,600 | h pS87 | 14,540 |
| h 1-101 | 10,062 | h 65-140 | 7,872 | h pY125 | 14,540 |
| h 1-110 | 11,045 | h 71-140 | 7,374 | h pY125/S129 | 14,620 |
| h 1-114 | 11,457 | h Δ111-115 | 13,933 | h pS129 | 14,540 |
| h 1-115 | 11,572 | h Δ111-115/133-135 | 13,526 | h pY125/pY133/pY136 | 14,745 |
| h 1-119 | 12,015 | h Δ120-125 | 13,770 | m 1-120 | 12,222 |
| h 1-120 | 12,112 | h Δ133-135 | 14,054 | m Δ120-125 | 13,881 |
| h 1-122 | 12,341 | h FL | 14,460 | m Δ133-135 | 14,079 |
| h 1-123 | 12,470 | h Y39F | 14,444 | m 20-140 | 12,551 |
| h 1-124 | 12,541 | h Y133/136F nY39 | 14,490 | m FL | 14,485 |
| h 1-133 | 13,627 | h nY39 | 14,522 | m E114A | 14,427 |
| h 1-135 | 13,871 | h nY125 | 14,522 | m D115A | 14,441 |
| h 5-140 | 13,968 | h pY39 | 14,540 | m pS129 | 14,580 |

aSyn = alpha-synuclein; Da = dalton; FL = full-length; h = human; m = mouse; Mw = molecular weight

**Supplementary Table 3:** A list of all the purified antibodies and their epitopes after finalisation of the monoclonal antibody generation programmes.

| antibody full name | antibody short name | epitope | species/ clonality | isotype | reactivity | concentration (mg/mL) | amount (mg) |
|---|---|---|---|---|---|---|---|
| LASH-EGT403 6D10-F10 | LASH-EGT403 | 1-5 | mus mc | IgG1 K | hu, mus | 1.1 | 56.2 |
| LASH-EGT410 5B10-A12 | 5B10-A12 | 1-10 | mus mc | IgG1 K | hu, mus | 1.4 | 56.2 |
| LASH-EGT406 2F10-E12 | 2F10-E12 | 110-115 | mus mc | IgG1 K | hu, mus | 1.8 | 79.7 |
| LASH-EGT405 7H10-E12 | 7H10-E12 | 115-125 | mus mc | IgG1 K | hu | 1.6 | 38.4 |
| LASH-EGT408 4E9-C12 | 4E9-C12 | 121-132 | mus mc | IgG1 K | hu | 1.2 | 35.5 |
| LASH-EGT405 2C4-B12 | 2C4-B12 | 123-125 | mus mc | IgG1 K | hu | 1.0 | 28.7 |
| LASH-EGT408 4E9-G10 | 4E9-G10 | 120-125 | mus mc | IgG1 K | hu | 0.9 | 26.5 |
| LASH-EGT410 6B2-D12 | 6B2-D12 | 126-132 | mus mc | IgG1 K | hu, mus | 1.2 | 37.2 |
| LASH-EGT410 1F10-B12 | 1F10-B12 | 121-125 | mus mc | IgG1 K | hu | 1.2 | 36.7 |
| LASH-EGT406 6A3-E9 | 6A3-E9 | aSyn-120 | mus mc | IgG1 K | hu | 1.0 | 23.0 |
| LASH-EGT416 5E1-G8 | 5E1-G8 | nY39 | mus mc | IgG1 K | hu, mus | 1.3 | 52.2 |
| LASH-EGT416 5E1-C10 | 5E1-C10 | nY39 | mus mc | IgG1 K | hu, mus | 1.3 | 53.1 |

aSyn = alpha-synuclein; hu = human; IgG = immunoglobulin G; K = kappa; mc = monoclonal; mus = mouse

**Supplementary Table 4:** A list of all the primary and secondary antibodies used in this study.

| primary antibodies | | | | | | |
|---|---|---|---|---|---|---|
| **antibody name** | **epitope** | **species/ clonality** | **reactivity** | **concentration (mg/mL)** | **company** | **catalogue #** |
| LASH-EGT403 | aSyn 1-5 | mus mc | h, m | 1.12 | - | - |
| 5B10-A12 | aSyn 1-10 | mus mc | h, m | 1.40 | - | - |
| LASH-EGTNter | aSyn 1-20 | rab pc | h, m | 1.34 | - | - |
| LASH-BL 34-45 | aSyn 34-45 | mus mc | h, m | 1.00 | Biolegend | 849102 |
| LASH-BL 80-96 | aSyn 80-96 | mus mc | h, m | 1.00 | Biolegend | 848302 |
| BD SYN-1 | aSyn 91-99 | mus mc | h, m | 0.25 | BD Transduction | BD610787 |
| BL 4B12 | aSyn 103-108 | mus mc | h | 1.00 | Biolegend | 807801 |
| 2F10-E12 | aSyn 110-115 | mus mc | h, m | 1.81 | - | - |
| AB LB509 | aSyn 115-122 | mus mc | h | 1.00 | Abcam | ab27766 |
| 7H10-E12 | aSyn 115-125 | mus mc | h | 1.60 | - | - |
| LASH-BL 117-122 | aSyn 117-122 | mus mc | h | 1.53 | Biolegend | 848601 |
| LASH-BL A15127A | aSyn 120-122 | mus mc | h | 5.19 | Biolegend | 848401 |
| 4E9-G10 | aSyn 120-125 | mus mc | h | 0.89 | - | - |
| 1F10-B12 | aSyn 121-125 | mus mc | h | 1.23 | - | - |
| 4E9-C12 | aSyn 121-132 | mus mc | h | 1.18 | - | - |
| 2C4-B12 | aSyn 123-125 | mus mc | h | 0.96 | - | - |
| 6B2-D12 | aSyn 126-132 | mus mc | h, m | 1.24 | - | - |
| AB 134-138 | aSyn 134-138 | rab pc | h, m | 1.00 | Abcam | ab131508 |
| LASH-BL pY39 | aSyn pY39 | mus mc | h | 1.00 | Biolegend | 849201 |
| LASH-EGT pY39 | aSyn pY39 | rab pc | h, m | 0.50 | - | - |
| LASH pS87 | aSyn pS87 | rab pc | h | 0.40 | - | - |
| LASH-EGT pY125 | aSyn pY125 | rab pc | h | 0.20 | - | - |
| AB pY125 | aSyn pY125 | rab pc | h | 0.50 | Abcam | ab10789 |
| AB EP1536Y | aSyn pS129 | rab mc | h, m | 2.68 | Abcam | ab51253 |
| LASH-EGT pS129 | aSyn pS129 | rab pc | h, m | 0.10 | - | - |
| BL 81A | aSyn pS129 | mus mc | h, m | 1.00 | Biolegend | 825701 |
| BL 81A biotin | aSyn pS129 | mus mc | h, m | 0.50 | Biolegend | 824704 |
| AB MJF-R13 | aSyn pS129 | rab mc | h, m | 4.20 | Abcam | ab168381 |
| AB pY133 | aSyn pY133 | rab pc | h, m | 1.50 | Abcam | ab194910 |
| AB pY136 | aSyn pY136 | rab pc | h, m | 1.00 | Abcam | ab131491 |
| LASH-EGT nY39 | aSyn nY39 | rab pc | h, m | 0.53 | - | - |
| 5E1-G8 | aSyn nY39 | mus mc | h, m | 1.31 | - | - |
| 5E1-C10 | aSyn nY39 | mus mc | h, m | 1.27 | - | - |
| 6A3-E9 | aSyn-120 | mus mc | h | 0.96 | - | - |
| AB β-actin AC-15 | β-actin | mus mc | h, m | 1.00 | Abcam | ab6276 |
| AB anti-MAP2 | MAP2 | ch pc | m, r | na | Abcam | ab5392 |
| TF AT8 | tau pS202/T205 | mus mc | h, m, r | 0.20 | ThermoFisher | MN1020 |
| Agilent 6F/3D | a-beta | mus mc | h | na | Agilent | M0872 |
| LSBio 2E2-D3 | TDP-43 | mus mc | h | 0.48 | LSBio | LS-B4521 |

| secondary antibodies | | | |
|---|---|---|---|
| **antibody name** | **concentration (mg/mL)** | **company** | **catalogue #** |
| DAPI 461 | 2.00 | ThermoFisher | D1306 |
| goat anti-mouse Alexa Fluor 488 | 2.00 | ThermoFisher | A-11029 |
| donkey anti-rabbit Alexa Fluor 488 | 2.00 | ThermoFisher | A-21206 |
| goat anti-chicken Alexa Fluor 568 | 2.00 | ThermoFisher | A-11041 |
| donkey anti-rabbit Alexa Fluor 568 | 2.00 | ThermoFisher | A-10042 |
| donkey anti-mouse Alexa Fluor 647 | 2.00 | ThermoFisher | A-31571 |
| donkey anti-rabbit Alexa Fluor 647 | 2.00 | ThermoFisher | A-31573 |
| IRDye 680RD goat anti-mouse | - | Li-Cor | 926-68070 |
| IRDye 800CW goat anti-rabbit | - | Li-Cor | 926-32211 |

a-beta = amyloid-beta; aSyn = alpha-synuclein; ch = chicken; h = human; MAP2 = microtubule-associated protein 2; mc = monoclonal; mus or m = mouse; pc = polyclonal; rab or r = rabbit; tau = tubulin-associated unit; TDP-43 = transactive response DNA-binding protein 43kDa

**Supplementary Table 5:** Optimised IHC settings for the antibodies used on human post-mortem brain tissues.

| antibody | epitope | IHC: dilution | IHC: antigen retrieval | IF: dilution* |
|---|---|---|---|---|
| LASH-EGT403 | aSyn 1-5 | 1:50 | FA | na |
| 5B10-A12 | aSyn 1-10 | 1:5,000 | AC+FA | na |
| LASH-EGTNter | aSyn 1-20 | 1:15,000 | AC+FA | 1:2,000 |
| LASH-BL 34-45 | aSyn 34-45 | 1:30,000 | FA | 1:10,000 |
| LASH-BL 80-96 | aSyn 80-96 | 1:20,000 | FA | na |
| BD SYN-1 | aSyn 91-99 | 1:5,000 | FA | na |
| BL 4B12 | aSyn 103-108 | 1:100,000 | AC+FA | 1:5,000 |
| 2F10-E12 | aSyn 110-115 | 1:10,000 | FA | 1:500 |
| AB LB509 | aSyn 115-122 | 1:25,000 | AC+FA | na |
| 7H10-E12 | aSyn 115-125 | 1:9,000 | AC+FA | na |
| LASH-BL 117-122 | aSyn 117-122 | 1:80,000 | AC+FA | na |
| LASH-BL A15127A | aSyn 120-122 | 1:15,000 | AC+FA | na |
| 4E9-G10 | aSyn 120-125 | 1:4,000 | AC+FA | na |
| 1F10-B12 | aSyn 121-125 | 1:15,000 | AC+FA | na |
| 4E9-C12 | aSyn 121-132 | 1:10,000 | AC+FA | na |
| 2C4-B12 | aSyn 123-125 | 1:12,000 | AC+FA | na |
| 6B2-D12 | aSyn 126-132 | 1:500 | FA | na |
| AB 134-138 | aSyn 134-138 | 1:25,000 | AC+FA | 1:100 |
| LASH-BL pY39 | aSyn pY39 | 1:2,000 | AC+FA | na |
| LASH-EGT pY39 | aSyn pY39 | 1:500 | AC+FA | 1:50 |
| LASH pS87 | aSyn pS87 | 1:600 | FA | na |
| LASH-EGT pY125 | aSyn pY125 | 1:100 | FA | na |
| AB pY125 | aSyn pY125 | 1:500 | FA | na |
| AB EP1536Y | aSyn pS129 | 1:60,000 | AC+FA | na |
| LASH-EGT pS129 | aSyn pS129 | 1:200 | FA | na |
| AB pY133 | aSyn pY133 | 1:400 | AC+FA | na |
| AB pY136 | aSyn pY136 | 1:100 | FA | na |
| LASH-EGT nY39 | aSyn nY39 | 1:1,000 | FA | na |
| 5E1-G8 | aSyn nY39 | 1:4,000 | AC+FA | na |
| 5E1-C10 | aSyn nY39 | 1:250 | AC+FA | na |
| 6A3-E9 | aSyn-120 | 1:2,500 | AC+FA | na |
| BL 81A biotin | aSyn pS129 | na | na | 1:500 |
| TF AT8 | tau pS202/T205 | 1:600 | AC+FA | na |
| Agilent 6F/3D | a-beta | 1:100 | AC+FA | na |
| LSBio 2E2-D3 | TDP-43 | 1:8,000 | AC | na |

*AC+FA pre-treatment was applied in IF studies for all antibodies. a-beta = amyloid-beta; AC = autoclave; aSyn = alpha-synuclein; FA = formic acid; IF = immunofluorescence; IHC = immunohistochemistry; na = not applicable; tau = tubulin associated unit; TDP-43 = transactive response DNA-binding protein 43kDa
